# Cell-type dependence of effects from transcranial electric brain stimulation

**DOI:** 10.64898/2026.04.19.719457

**Authors:** Susanne Dahle, Gaute T. Einevoll, Torbjørn V. Ness

## Abstract

There is an urgent need for better treatment options for many neurological conditions, including Alzheimer’s disease, Parkinson’s disease, depression, and epilepsy. Transcranial electrical stimulation (tES) is a non-invasive, safe, inexpensive, and promising method that could address some of this unmet need. The therapeutic value of tES has been well demonstrated, but the effect is highly variable. To enable tES to reach its full potential requires a better understanding of how tES modulates neural activity so that tES treatments can be tailored to specific neurological conditions and individual patients.

The neural response to tES is, however, highly complex, and the parameter space involved in optimizing tES treatments is daunting. This has made it difficult to obtain general insights into how tES modulates neural activity, and a central challenge lies in the cell-type-specific and frequency-dependent nature of these responses.

In this study, we investigate cell-type-specific neuronal responses to tES over a broad frequency range, using a large database of biophysically detailed neuron models. We find that pyramidal cells respond strongly to low-frequency tES, but their responses drop sharply with frequency. In contrast, inhibitory neurons show a smaller reduction and, on average, become more responsive than pyramidal cells above ∼ 60 Hz.

By leveraging a reciprocity theorem we demonstrate that the effect of tES on a given cell-type is proportional to the frequency-dependent current-dipole moment that determines the EEG-signal contribution of this cell-type. We further identified the dendritic asymmetry as key in determining tES responses across the frequency spectrum. Counterintuitively, we also found that while total cell length increases tES sensitivity at low frequencies, it can have the opposite effect at high frequencies. Furthermore, we derived an analytical formula for idealized neuron models which can approximately predict the tES sensitivity of different cell types at any given frequency.

By characterizing the role of morphology and stimulation frequency in determining tES responses of single cells, this is an important step towards a better understanding of tES at the fundamental level. These results also provide an efficient and accurate method for characterizing and comparing the tES responses of different neural populations across the frequency spectrum, which facilitates optimizing tES for cell-type specific targeting.

**Author summary:** 

## 1 Introduction

Neurological and psychiatric disorders impose a massive burden on those affected and on society as a whole [1], and there is an urgent need for improved interventions that can reduce disability and improve the quality of life for affected patients [2].

Transcranial electric stimulation (tES) is a promising non-invasive neuromodulation technique that has shown potential in treating various neurological conditions, including Alzheimer’s disease [3], Parkinson’s disease [4], depression [5], chronic pain [6], cognitive fatigue [7], stroke [5], and epilepsy [8]. By modulating neural activity non-invasively, tES offers a safe [6], low-cost, and portable intervention that can potentially address some of the unmet needs in neurological disease management [9].

tES is increasingly used both in clinical practice and in basic neuroscience research [9]. Despite their widespread application, the mechanisms by which externally applied electric fields influence neural circuits remain only partially understood [10, 11], and results are highly variable [9, 12]. A better understanding of tES could help reduce the therapeutic variability and pave the way for efficient personalized closed-loop treatments [9, 13, 14].

Biophysical modeling provides a powerful framework for investigating the mechanisms underlying tES [15–20]. Using the well-established cable-equation description of biophysically detailed neuron models combined with equally well-established volume-conductor theory for brain tissue [21], such computational models enable a mechanistic description of how externally applied fields modulate neuronal membrane potentials [11, 22–26].

Although the effect of a given tES configuration on a given cell model can, in principle, be simulated numerically based on well-established biophysics, it has proven challenging to gain a good understanding of how to optimize tES treatments in practice. One reason for this is the vast parameter space [9], making it hard to extract qualitative insight into how different types of tES affect different types of neurons and neural elements. It is, for example, well established that electric fields interact differently with distinct neuronal cell types due to their unique morphologies and membrane properties [27, 28]. Several studies have used databases of biophysically detailed cell models to numerically evaluate how different cell types react to low-frequency (≤ 100 Hz) tES [19, 20, 29, 30], but although these studies have yielded important insights, the exact relationship between cellular properties and tES sensitivity across a broad frequency spectrum has been difficult to extract.

A second reason that it is difficult to optimize tES treatments is the complexity of neural network dynamics in response to tES. In principle, the effect of tES can be simulated using large-scale networks of biophysically detailed cell models, however, this comes at a massive computational cost, making comprehensive parameter scans out of reach [31]. A more common way to investigate network effects is through point-neuron networks or mean-field models [18, 32, 33]. However, the effect of tES at the fundamental level stems from electric potential gradients over neural morphologies [26, 34, 35], and this level of detail is not represented in these approaches. Simplifying assumptions regarding how different neurons respond to tES have therefore been used, such as relying on neuron models with the minimum possible (two-compartment) spatial structure [32, 33], or incorporating tES as a current input given exclusively to excitatory cells [15, 18]. Although such simulations have proven helpful for simulating the network-level effect of tES, the potentially inaccurate representation of tES at the fundamental level of single cells is a substantial weakness. What is needed is therefore a principled understanding of cell-type- and frequency-specific effects of tES, which can be used in simplified network simulations based on point neurons and even mean-field models.

In this work, we take advantage of the extensive database of biophysically detailed neuron models provided by the Blue Brain Project [36] to explore how the effects of tES depend on cell type across a broad frequency spectrum. In line with previous studies, the different cell-types are found to have quite distinct frequency characteristics [19, 20, 29, 30]. For example, while the somatic membrane potentials of excitatory pyramidal neurons are on average observed to respond most strongly to low-frequency stimulation, inhibitory neurons are on average affected more strongly than pyramidal cells at higher frequencies, above 60 Hz or so.

Rather than just numerically quantifying the relationship between different cell types and susceptibility to tES, we apply a reciprocity theorem [37] to gain deeper insight into which neuronal features shape tES responses. This reveals that the effect of tES on the soma membrane potential of a neuron is directly given by the current dipole moment set up by injecting a current into the same neuron. Furthermore, an analytic formula is derived for the tES response of idealized neuron models with arbitrary numbers of dendrites.

Our results lead to a better, principled understanding of how tES affects different cell types—for both biophysically detailed and simplified cell models—across a broad frequency spectrum. Furthermore, we provide a highly efficient approach for accurately incorporating this effect into network simulations, in a way that facilitates parameter scans. This amounts to an important step towards the goal of maximizing the therapeutic efficiency and consistency of tES.

## 2 Results

### 2.1 Frequency response to electric stimulations is cell-type specific

To mimic transcranial electric stimulation (tES) we used a spatially uniform external electric field—the so-called quasi-uniform approximation—because the large and distant electrodes applying the current during tES create electric fields in the brain with low spatial gradient on the size scale of individual neurons [20, 35, 38]. We used an amplitude of 1 mV/mm, as this is a typical value that can be safely reached in human applications [6, 29, 35, 39], and focused on electric fields with frequencies ranging from 1 Hz to 2 kHz (see Methods). We mainly focused on fields oriented along the cortical depth axis along the apical dendrites of pyramidal cells (defined as the *z*-axis), as pyramidal cells are known to be most sensitive to electric fields with this orientation [10, 27, 33, 35, 40].

We simulated the somatic membrane potential (*V*_m_) response of the neocortical neurons from the Blue Brain Project [36] in response to this field. The extracellular stimulation was modeled using the standard approach, that is, through using the induced extracellular potentials directly outside each cellular compartment as boundary conditions (see Methods). The neurons were made passive by removing all active conductances because the low intensity current used during tES on humans is generally insufficient to directly induce action potentials [10, 11]. We confirmed that the active and passive cell models gave very similar results, although some models exhibited differences below about 10 Hz (Fig S1), due to the presence of stochastic potassium channels or *I*_h_ channels in some of active Blue Brain models [41–43]. In total, we considered 1035 different cell models with widely variable morphological features. We grouped neurons into three classes: Pyramidal cells (*N*=60), spiny stellate cells (*N*=5), and inhibitory neurons (*N*=970). Spiny stellate cells constitute a small subset of excitatory interneurons, but since the remaining interneurons are inhibitory, spiny stellate cells were analyzed separately.

There was a large variability in single-cell responses, but pyramidal cells generally had higher responses to low-frequency tES than interneurons and spiny stellate cells, with the notable exception of inhibitory L4 bipolar (BP) cells. This subclass had the strongest low-frequency tES response of all tested neurons (Fig 1B). Pyramidal-cell responses decreased markedly with frequency, while most inhibitory neurons (again with the exception of the L4 BP cells) and spiny stellate cells had less pronounced frequency dependencies in their responses. From Fig 1C, it is not immediately apparent which cell types are most strongly affected by tES at higher frequencies, and the average somatic *V*_m_ response was calculated (Fig 1D). At stimulation frequencies above about 60 Hz interneurons had a higher average response than pyramidal cells (p-value = 0.0081 from one-sided Mann-Whitney U test conducted for *f* = 1000 Hz).

**Fig 1.**
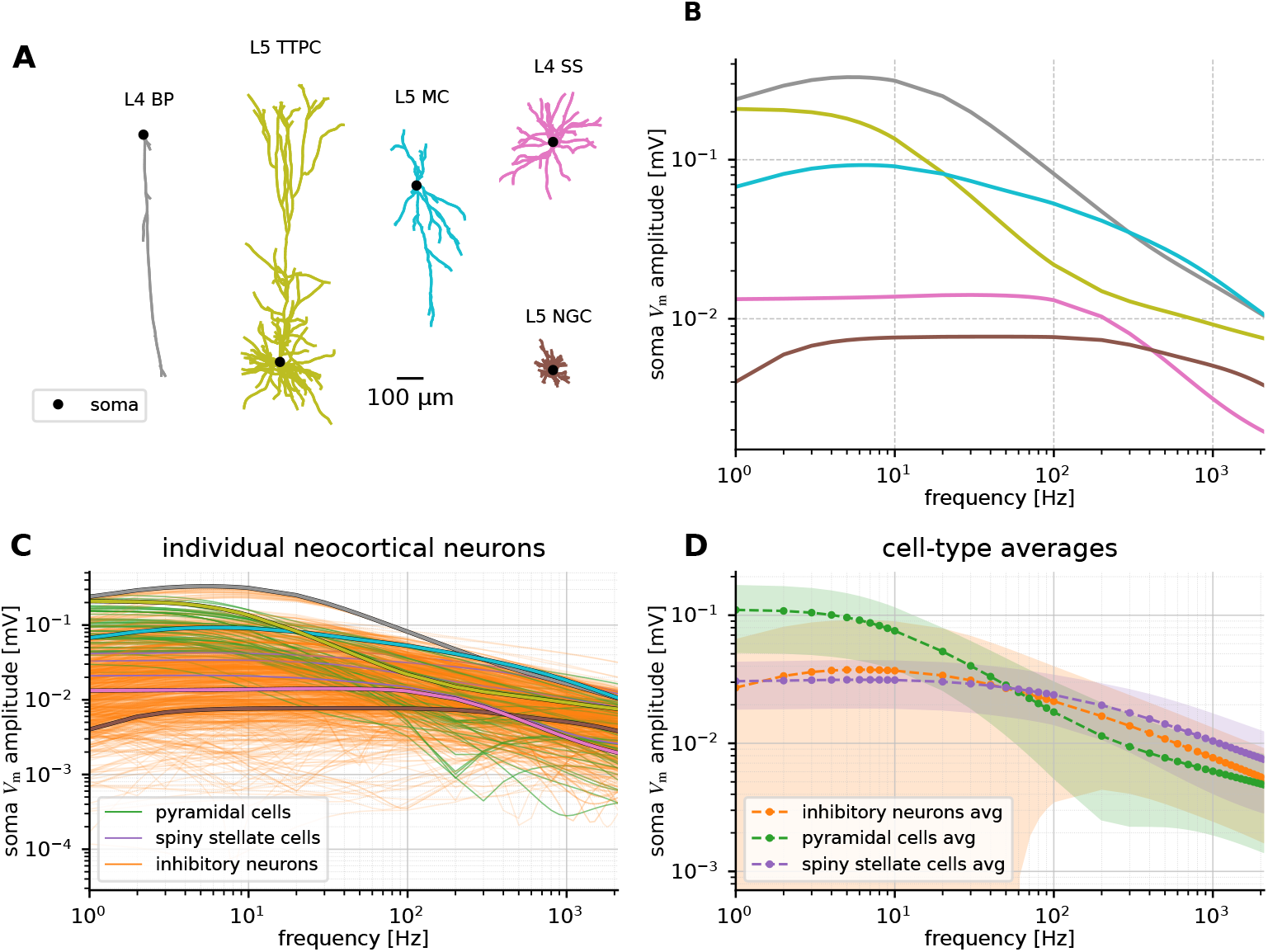
tES sensitivity is cell-type specific and frequency dependent. **A:** Morphologies of example cell models, including three inhibitory neurons (cortical layer 4 (L4) bipolar cell (BP), L5 Martinotti cell (MC), L5 neurogliaform cell (NGC)), one pyramidal cell (L5 thick-tufted pyramidal cell (TTPC)) and one L4 spiny stellate cell (SS). **B:** Somatic *V*_m_ amplitude response of the five example cell models to external electric field stimulation of 1 mV/mm in the *z*-direction, perpendicular to the cortical surface with frequencies ranging from 1 to 2000 Hz. **C:** All 1035 neocortical neuron models stimulated with an external electric field like in panel B. Example cells from panel B are marked. **D:** Average somatic *V*_m_ amplitude for the different cell types.

As demonstrated here and by previous studies [20, 27–30], neurons exhibit cell-type-specific responses to tES. The cell-type-specific responses are closely linked to differences in morphological properties [20, 27, 37]. Pyramidal cells, for example, are generally longer than interneurons in the *z*-direction perpendicular to the cortical surface (Fig 2A). This is caused by their long apical dendrite in the upwards direction (Fig 2B), while their spatial extent in the downward direction is more similar to inhibitory neurons (Fig 2C). This also imply that pyramidal cells are, on average, more asymmetric than interneurons (Fig 2D). Somatic and dendritic diameters are, however, relatively similar for pyramidal cells and interneurons (Fig 2E,F).

**Fig 2.**
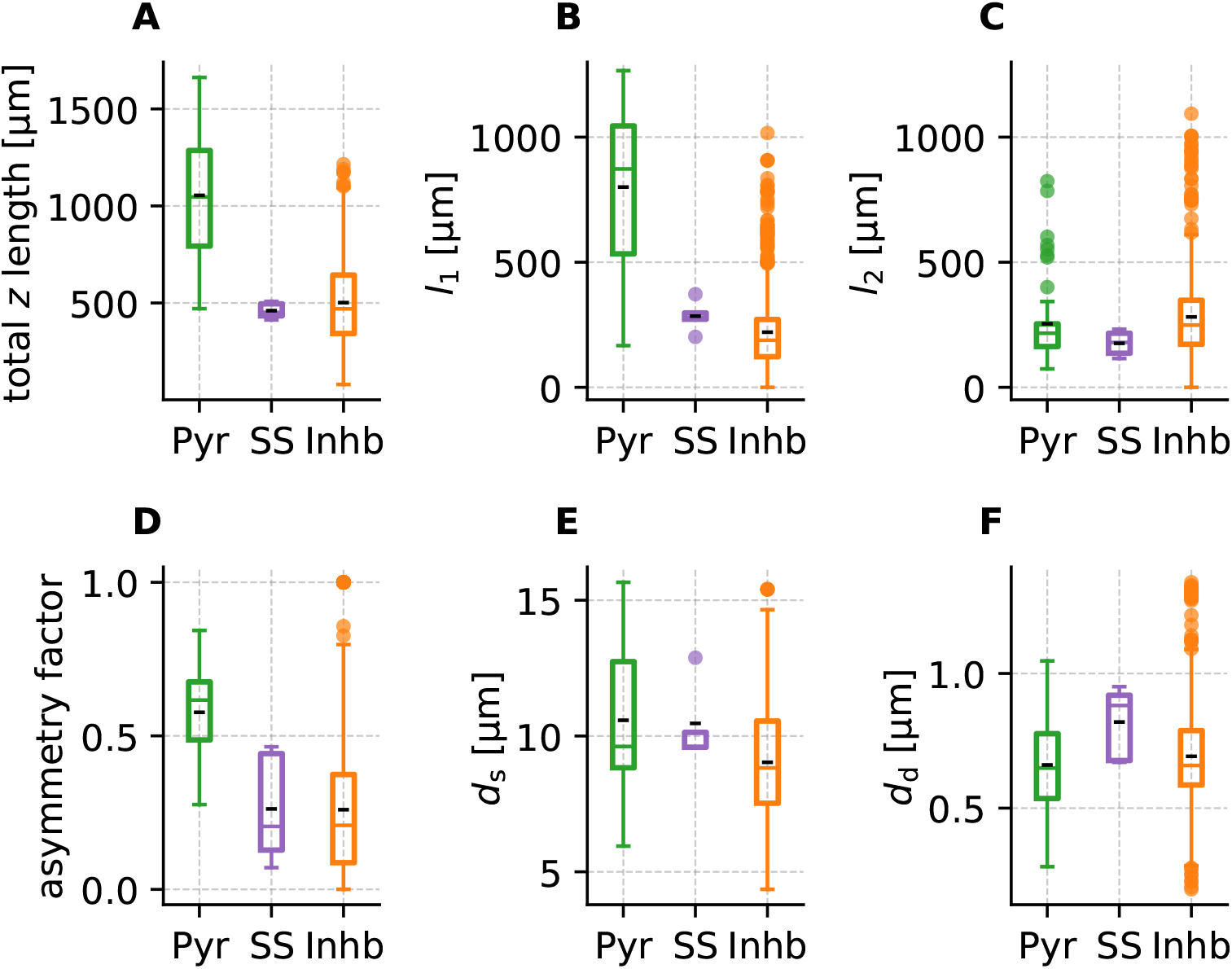
Morphological properties are highly cell-type specific. Box plots of different morphological properties for three classes of neurons, including pyramidal cells (Pyr), stellate cells (SS), and inhibitory interneurons (Inhb). **A:** The total length in the *z*-direction, calculated as the distance between the upper and lower endpoints. **B:** The length of the neuron above the soma (*l*_1_), i.e, the distance from the soma to the upper endpoint. **C:** The length of the neuron below the soma (*l*_2_), i.e, the distance from the soma to the lower endpoint. **D:** Asymmetry factor, defined as |*l*_1_ − *l*_2_ | */*(*l*_1_ + *l*_2_). **E:** Average diameter of the compartments pointing in *z*-direction. **F:** Diameter of the soma compartment. See Methods for more details about how the properties were found. Median marked with a colored line spanning the entire box, mean marked with a shorter black line. The boxes represent the interquartile (IQR) range (25th–75th percentile), containing the middle 50% of the data. Whiskers extend to 1.5 *×* IQR, and dots denote outliers beyond 1.5 *×* IQR.

The exact relationship between sensitivity to tES and morphology is, however, not *a priori* clear. To investigate how morphological properties influence somatic *V*_m_ amplitude responses to tES, we performed a linear regression analysis combined with a feature importance assessment to quantify each property’s contribution. The selected properties were soma diameter (*d*_s_), asymmetry factor, average diameter of compartments oriented in the *z*-direction (*d*_d_), and total *z*-directed length (Fig 3).

**Fig 3.**
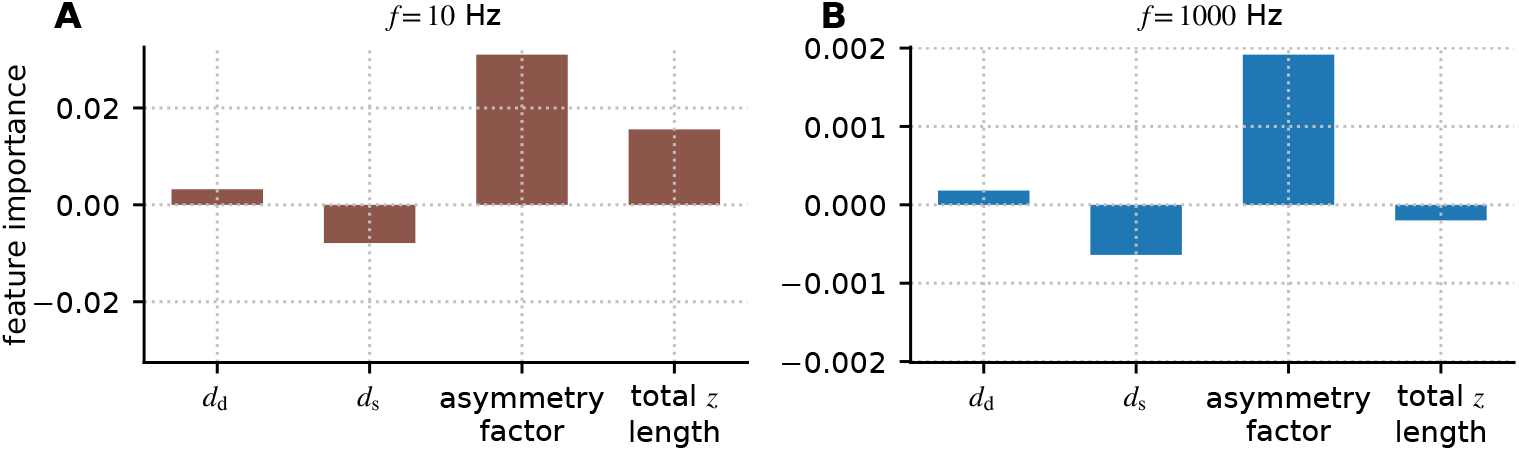
Feature importance analysis highlight the importance of the asymmetry factor, and indicate opposite effects of total cell length at low and high frequencies. Linear regression models using the average diameter of compartments oriented in the *z*-direction (*d*_d_), soma diameter (*d*_s_), asymmetry factor, and total *z*-directed length to predict soma *V*_m_ amplitudes induced by an electric field applied in the *z*-direction (see Methods for how the properties were determined). **A:** At *f* = 10 Hz, the regression model achieved *R*^2^ = 0.64. The asymmetry factor and total *z*-directed length have the highest feature importance values. The feature importance values are 0.0032 (*d*_d_), −0.0079 (*d*_s_), 0.031 (asymmetry factor) and 0.016 (total *z* length). **B:** At *f* = 1000 Hz, the regression model achieved *R*^2^ = 0.15. The asymmetry still has the highest feature importance value, while the total *z* length have a slight negative feature importance value. The feature importance values are 0.00018 (*d*_d_), − 0.00064 (*d*_s_), 0.0019 (asymmetry factor) and − 0.00020 (total *z* length). The feature importance values are the regression coefficients, meaning the model’s predicted change in soma amplitude (in mV) per 1 standard deviation increase in that predictor. To minimize multicollinearity, we confirmed that the selected morphological properties were at most weakly correlated (Fig S2). Feature importance analysis was also performed using idealized ball-and-two-sticks neuron models where the morphological features were fully independent, giving similar results (Fig S8).

The linear regression model achieved *R*^2^ = 0.64 at *f* = 10 Hz, indicating that a substantial proportion of the variance in somatic *V*_m_ amplitude could be explained by the selected morphological properties. In contrast, at *f* = 1000 Hz, the model performance dropped markedly to *R*^2^ = 0.15, demonstrating limited predictive power at high frequencies. The limited predictive power of each individual morphological feature was also confirmed by single-feature regression models, indicating a low correlation between any individual morphological property and the somatic *V*_m_ response (Fig S3). Through visual inspection of the relationship between each morphological property and somatic *V*_m_ response, it is also clear that the low *R*^2^ value is not due to non-linear relationships that are not captured by the regression model, but rather highly variable data (Fig S3).

The feature importance analysis revealed distinct frequency-dependent differences in how morphology contributes to tES responses (Fig 3). At 10 Hz, the asymmetry factor and total *z* length were the strongest contributors, with feature importance values of 0.031 and 0.016, respectively. These positive values indicate that increased asymmetry and greater total *z*-length are associated with larger soma *V*_m_ amplitudes at low frequencies.

At 1000 Hz, all feature importance values were substantially reduced in magnitude, consistent with the low *R*^2^ value. The asymmetry factor remained the largest contributor (0.0019), although its effect size was markedly diminished compared to 10 Hz. In contrast, the contribution of total *z*-directed length dropped dramatically and became slightly negative (-0.00020), suggesting that increased neuronal length is weakly associated with a reduced tES response.

Together, these results indicate that morphological properties shape the neuron’s responses to tES, but the high variability in tES responses and morphological properties makes the exact link non-trivial to extract. At low stimulation frequencies, particularly asymmetry and longitudinal extent are important, which is consistent with previous findings [20, 27, 29, 30, 37]. However, at high frequencies, the relationship between morphology and response to tES becomes even less clear cut.

### 2.2 Applying the reciprocity theorem in the context of tES

The tES response of different cell types varies in both amplitude and frequency dependence, and it is not trivial to link tES responses directly to differences in morphological properties, such as spatial extent, dendritic and somatic diameters, or branching patterns. It was, however, recently demonstrated that the reciprocity theorem can be helpful for analyzing the responses of neurons to tES [37]. In the context of tES, the reciprocity theorem states that the somatic membrane potential response 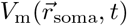 resulting from an extracranial current source *I*_in_(*t*) at location 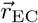 is, for a passive model, identical to the electroencephalography (EEG) signal 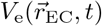 at location 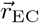 resulting from a somatic current input *I*_in_(*t*).

Note that typical current amplitudes used for tES (∼ mA) and somatic input (∼ pA) are vastly different, however, since passive cell models are linear, we are free to scale the current amplitudes and the resulting potentials. This approach has been demonstrated to be accurate even for active cell models, as long as the resulting membrane potentials are within the subthreshold regime [37], as expected for tES (with the exception of electro-convulsive therapy, where extremely high-amplitude currents are used [44]).

The reciprocity theorem can in the present context be mathematically expressed as

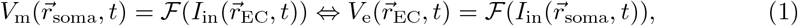

where ℱ is a function linking the stimulus current to the potential response [21]. The symbol ⇔ here indicate that the two sides are equivalent by the reciprocity theorem. The EEG-signal contribution from a neuron is directly proportional to the neuron’s current-dipole moment 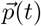 [21, 45],

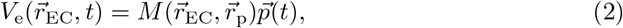

where *M* is the head model linking a current dipole 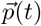 at a given location 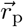 with the resulting EEG signal *V*_e_ at location 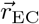.

The head model *M* can be anatomically and electrically detailed [46] but is often pre-solved through numerical methods like the finite element method and expressed in the form of a lead-field matrix linking a specific set of EEG scalp locations with a specific set of possible dipole locations in the brain [21, 47]. Because of radial symmetry around the axis of the cortical depth direction (*z*-axis), it is generally expected that only the dipole component oriented along the cortical depth direction, *p*_*z*_(*t*), contributes to the resulting EEG signal [45]. For a specific combination of a tES electrode location 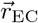 and cortical location 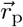. *M* reduces to a fixed scalar, here denoted *k*.

If *p*_*z*_(*t*) was calculated from a somatic input current with amplitude *I*_0_, and we are interested in the somatic response of the neuron to tES with an amplitude 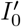, we have

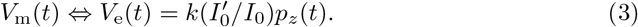

Note that *k* only depends on the head model, and will be the same for all cell models. Thus, we can study the susceptibility of neurons to tES—through their somatic membrane potential response—by calculating their current-dipole moments from somatic current input.

As independent verification and demonstration of this use of the reciprocity theorem, the five example neurons from Fig 1 were stimulated with somatic white noise current (*I*_in_), containing frequency components ranging from 1 to 2020 Hz, where each frequency component had an amplitude of 5 pA. The amplitudes of the *z*-component of the dipole moment (*p*_*z*_) were computed across stimulation frequencies and compared to soma *V*_m_ amplitudes from extracellular electric field stimulation. For all five models, *p*_*z*_ amplitudes followed the same frequency-dependent trend as the soma *V*_m_ amplitudes (Fig 4A-B).

**Fig 4.**
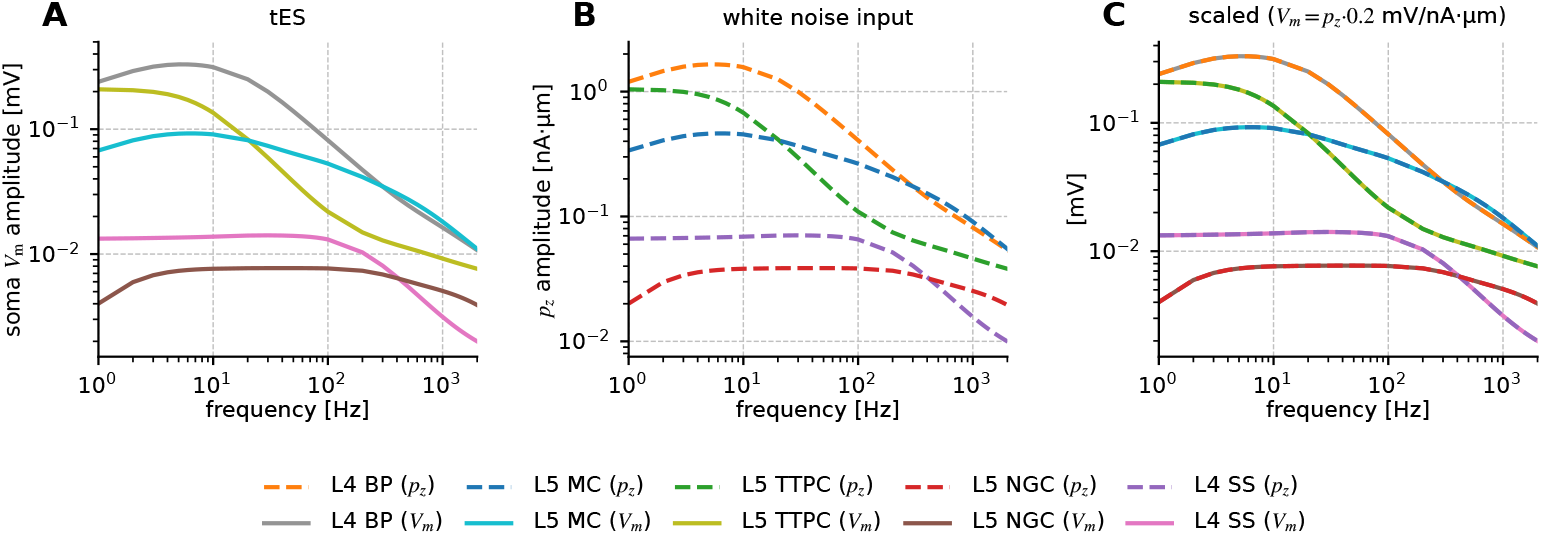
The reciprocity theorem can be used to calculate somatic membrane potential responses from tES. **A:** Somatic membrane potential (*V*_m_) amplitude in response to *z*-oriented tES across frequencies. **B:** *p*_*z*_ amplitudes across frequency in response to somatic white noise current input. **C:** Somatic *V*_m_ amplitudes (solid) and *p*_*z*_ amplitudes scaled with 0.2 mV/nAµm (dashed) in the same plot.

To directly compare membrane potential responses, we used the New York head model [47] and the same combination of tES and dipole locations as Ness et al. (2025) [37, their figure 6], resulting in *k* = 0.21 · 10^−9^ mV/nAµm. At this location, the electric field was 0.21 mV/mm per mA of applied tES current. A field amplitude of 1 mV/mm (like used here) would therefore require a current amplitude of 4.76 mA. The different current amplitudes result in a scaling factor of 4.76 mA/5 pA = 9.52 · 10^8^. Therefore, *V*_m_ = 0.21 · 10^−9^ · 9.52 · 10^8^ · *p*_*z*_ mV/nAµm= 0.2 · *p*_*z*_ mV/nAµm.

As shown in Fig 4C, the reciprocity-based *V*_m_ calculation gave indistinguishable results to those of the traditional approach to simulating the effect of tES, confirming that *p*_*z*_ can be used to study tES. In the following, we will leverage this to study the effects of neuronal morphology on the response to tES.

### 2.3 tES responses explained through reciprocity-based analysis

We have demonstrated that we can directly predict the somatic *V*_m_ response of a neuron to tES from the current dipole moment *p*_*z*_ resulting from somatic current input. In other words, the susceptibility of a neuron to tES is directly proportional to its dipole moment for somatic current input. This can be leveraged to illuminate the role of morphology and stimulation frequency in determining tES responses.

We therefore review some basic features of how *p*_*z*_ is shaped by cellular properties: *p*_*z*_ is determined by the distribution of membrane currents in the *z*-direction, where the somatic input current counts as a membrane current [37]. For a neuron with *N* compartments,

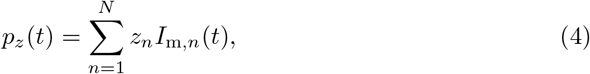

where *z*_*n*_ is the *z*-coordinate of compartment *n*, and *I*_m,*n*_ is the membrane current of the same compartment [48, 49].

From theory, we know that *p*_*z*_ will be maximized for spatially asymmetric current distributions, that is, when return currents are primarily either above or below the soma [21, ch. 8]. This implies an important role for the asymmetry of neural morphologies around the soma. *p*_*z*_ also increases with the separation between the somatic input current and the return currents, and long neurons are, therefore, more capable of generating large *p*_*z*_. Furthermore, thick dendrites will, in general, result in more distant return currents, while large somas will have the opposite effect and result in larger somatic return currents [37].

Because of intrinsic dendritic filtering [21, 48], the spatial distribution of membrane currents changes with frequency. We investigated this via the frequency-dependent length constant *λ*_AC_. Pettersen and Einevoll (2008) [50] defined *λ*_AC_ as the mean of the absolute value of the membrane current amplitude, weighted by distance. In essence, we can think of *λ*_AC_(*f*) as representing the average distance from the input site to the return currents at a given frequency [21], and it is therefore tightly linked to *p*_*z*_.

With complex notation (bold), the membrane current density for a single frequency component *f* is given by 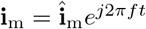. For a finite stick with length *l, λ*_AC_ can then be written as [50]

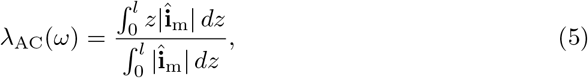

which for an infinite stick simplifies to

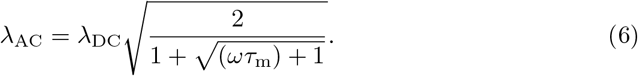

Here, *τ*_m_ = *R*_m_*C*_m_ is the membrane time constant, and *ω* = 2*πf* is the angular frequency.

We calculated *λ*_AC_ for neural sticks of different lengths using Eq (5) and Eq (6) to demonstrate how the distribution of return currents depends on both frequency and the length of the stick. The longer sticks had much higher *λ*_AC_, that is, more distant return-currents, than the shorter sticks at low frequencies, but they also had a much sharper decline in *λ*_AC_ with frequency (Fig 5). This can explain why pyramidal cells, on average, had more frequency-dependent responses to tES (Fig 1). However, *λ*_AC_ for the longer neurons never became lower than those of the shorter neurons. The change in *λ*_AC_ is, therefore, not sufficient to explain why interneurons had a higher tES response than pyramidal cells at high frequencies (Fig 1D). We therefore turned to the role of morphological asymmetry.

**Fig 5.**
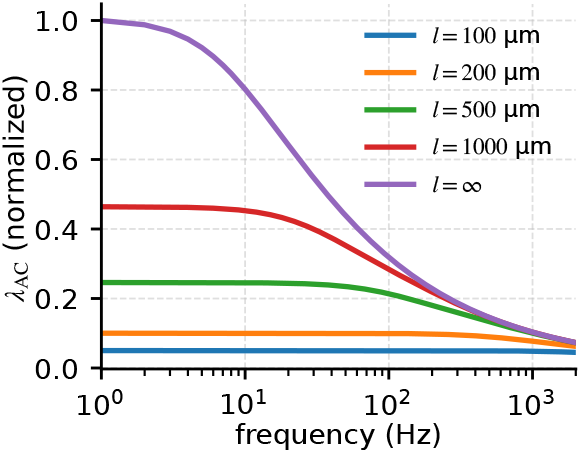
Longer neurites have more frequency dependent responses than shorter neurites. Frequency-dependent length constant as a function of frequency. All sticks had a diameter of 2 µm, with axial resistance *R*_i_ =150 Ωcm, membrane resistance *R*_m_ = 30000 Ωcm^2^, and membrane capacitance *c*_m_=1 µF/cm^2^. Figure adapted from Halnes et al. (2024) [21, p.63].

The current dipole moment of a cell resulting from somatic current input is highly dependent on the asymmetry of the morphology around the soma: *p*_*z*_ is maximized if all return currents are on one side of the soma, while it is zero in the limit of a perfectly symmetric distribution. To investigate how the reduction in *λ*_AC_ affects the asymmetry of membrane currents, *λ*_AC_ was found numerically above and below the soma for both idealized “ball-and-two-sticks” neurons and morphologically detailed neurons. The compartments of the neuron were separated into those above (*N*_above_) and below (*N*_below_) the soma, and the amplitudes of *I*_m_ in all compartments were used to calculate *λ*_AC_ as

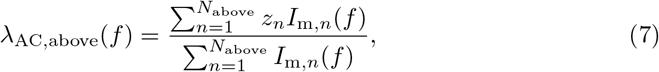

and

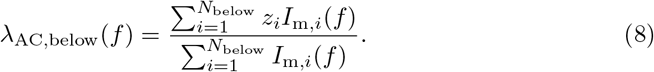

Eq (7) and (8) are similar to the definition of the frequency-dependent length constant *λ*_AC_ in Eq (5), except that the amplitude of the current is used instead of current density, and the neuron is discretized into compartments to approximate the solution numerically.

Note that the numerator in Eq (7) and Eq (8) for *λ*_AC_, is similar to *p*_*z*_ in Eq (4). However, with alternating current input, the phases of the membrane currents can vary for different compartments, while Eq (7) and Eq (8) for *λ*_AC_ only uses the amplitude. Therefore, while *λ*_AC_ is similar to *p*_*z*_, it specifically illustrates the distribution of membrane currents at different frequencies for different neuronal morphologies. We also separated *p*_*z*_ into *p*_*z*,above_ and *p*_*z*,below_, that is, the current-dipole moment calculated from all compartments above or below the soma, respectively. *p*_*z*_—which we use to measure the sensitivity of a cell to tES—is the sum of these two components.

For an example long ball-and-two-sticks model, the membrane current distribution was strongly frequency dependent (Fig 6B-C). At low frequencies, the return currents were biased towards the longest branch representing the apical dendrite (Fig 6C). This asymmetry resulted in a strong low-frequency net *p*_*z*_ (Fig 6C). At higher frequencies, *λ*_AC_ was, however, much smaller than the branch length both above and below the soma (Fig 6C), effectively making the cell highly symmetric. This resulted in a weak total *p*_*z*_ at high frequencies. The total current dipole moment was reduced from 0.80 nAµm at 10 Hz to 0.0012 nAµm at 1000 Hz, that is, a factor 670 smaller.

**Fig 6.**
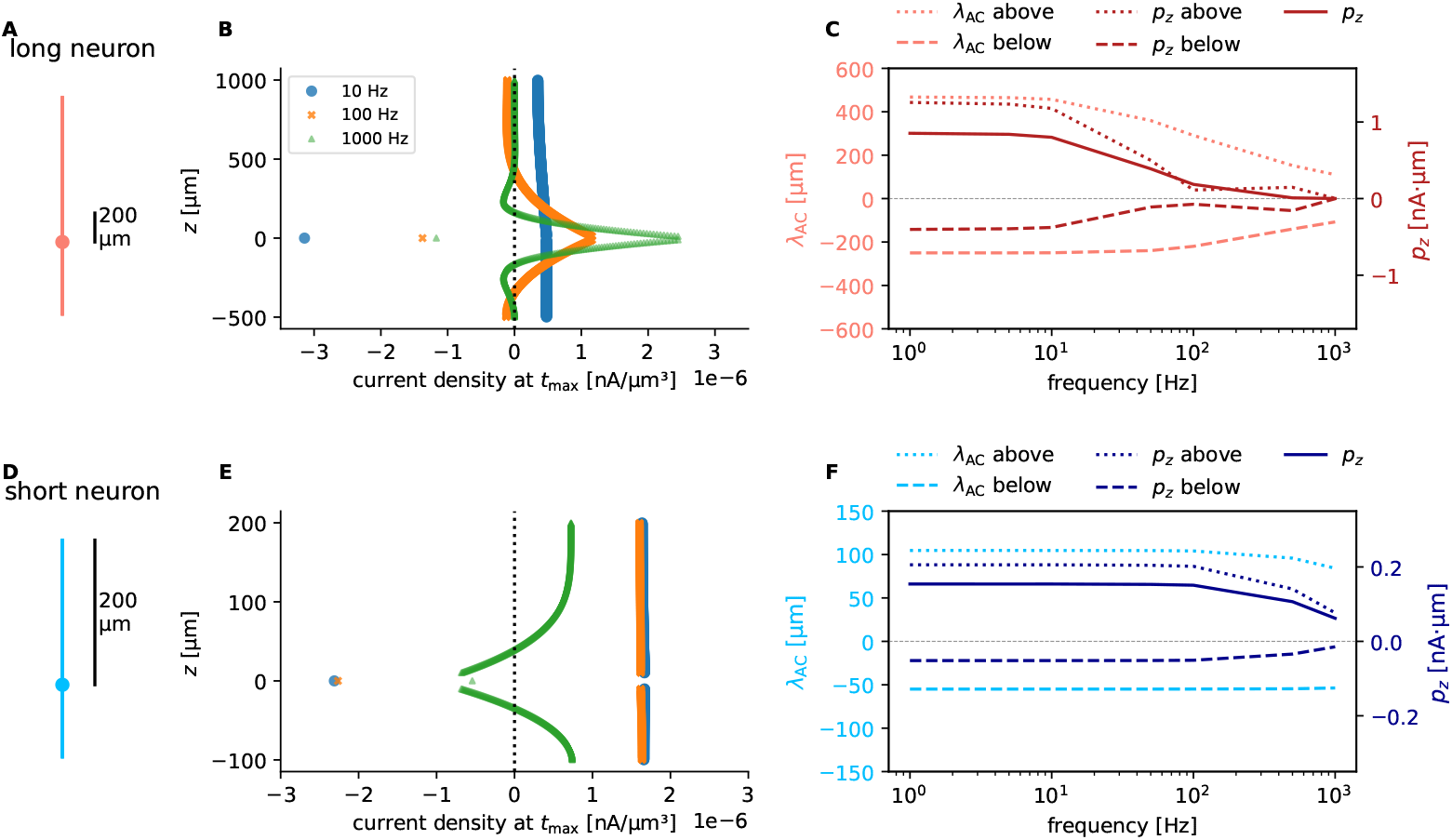
Long neurons can effectively become very symmetric at high frequencies. **A:** Length of the two sticks for the two different ball-and-two-sticks neurons. White-noise current is injected at the soma (dot). **B:** Resulting membrane current values for each compartment at different frequencies of the input current. The membrane current in each compartment was reconstructed as a sinusoidal signal using the amplitude and phase found via Fourier analysis (see Methods). The value is plotted for time *t*_max_, where *p*_*z*_ has its maximum value (amplitude). Note that membrane current in the soma is the injected current minus return current exiting the soma, so a lower *I*_m_ at *z* = 0 for *f* = 1000 Hz indicates more current exiting directly through the soma. **C:** Weighted average position of the amplitude of membrane currents above and below soma (*λ*_AC_), and the *p*_*z*_ value, *p*_*z*_ above and *p*_*z*_ below at *t*_max_. Except for the dendritic lengths, all parameters were the same: Dendritic diameters were 2 µm, somatic diameters were 20 µm, axial resistance was *R*_i_ =150 Ωcm, membrane resistance was *R*_m_ = 30000 Ωcm^2^, and membrane capacitance was *c*_m_=1 µF/cm^2^.

For an example short neuron, the spatial current distribution and *λ*_AC_ were less affected by frequency (Fig 6E-F), indicating that the dendritic branches were short enough for intrinsic dendritic filtering to have a limited effect on the spatial membrane current distribution. For the short neuron model the total current dipole moment was reduced from 0.15 nAµm at 10 Hz to 0.060 nAµm at 1000 Hz, that is, a reduction of only a factor 2.5. In contrast to the long neuron model, the short neuron model effectively retained its asymmetry at higher frequencies, resulting in the observed larger high-frequency *p*_*z*_ for the short neuron than for the long neuron.

The example long and short ball-and-two-sticks models had the same dendritic asymmetry. However, the membrane current asymmetry had very different frequency-dynamics in the long and short models. The asymmetry used in the feature importance analysis (Fig 3) was the dendritic asymmetry and not the frequency dependent membrane current asymmetry. The simplified models demonstrate that these two measures can diverge as frequency increases. Based on the observed behavior of the simplified models, we hypothesized that for low-frequency tES, pyramidal cells are, on average, more sensitive than interneurons due to their larger and more asymmetric morphologies. For high-frequency tES, however, pyramidal cells were hypothesized to be less sensitive than interneurons because, at such frequencies, they would be effectively more symmetric than the interneurons.

To investigate this mechanism in morphologically detailed neuron models, we again considered the five example neurons from Fig 1. The two example models with the most pronounced asymmetry around the soma are the L4 bipolar cell and the L5 pyramidal cell (Fig 7A). These two cell models also had the highest low-frequency *p*_*z*_ amplitudes (1.20 nAµm and 1.04 nAµm for the L4 BP and L5 PC, respectively, at 1 Hz). For the pyramidal cell, *p*_*z*,above_ was highly frequency-dependent, while *p*_*z*,below_ was close to frequency-independent (Fig 7C). At low frequencies, *p*_*z*,above_ dominated, while they were close to equal but opposite at high frequencies, resulting in a weak net *p*_*z*_.

**Fig 7.**
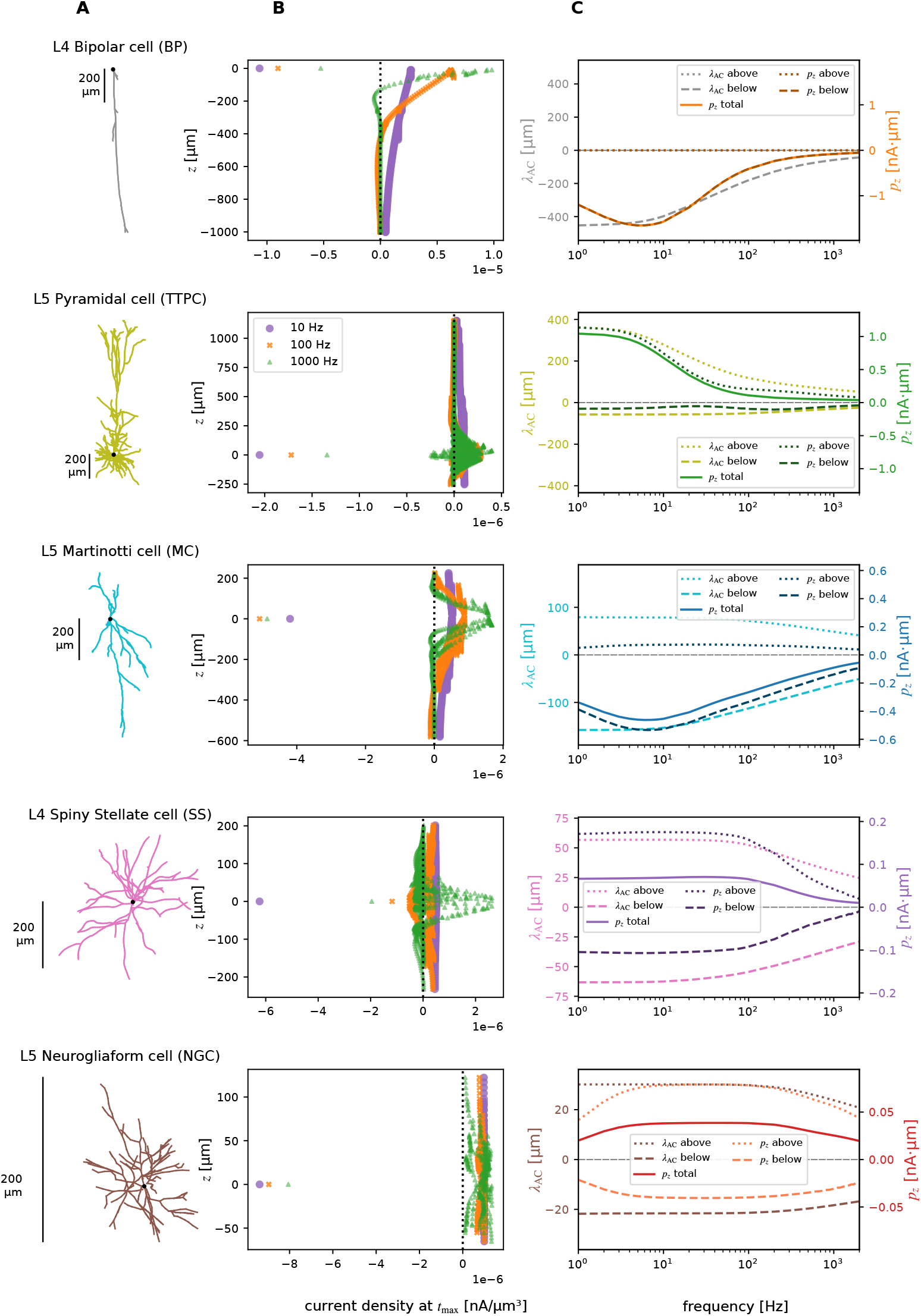
Effective asymmetry is frequency dependent and variable in neocortical neurons. **A:** Morphology of the neurons. **B:** Membrane current distribution at different frequencies of the input current. The membrane currents are plotted for time *t*_max_, where *p*_*z*_ has its maximum amplitude. Note, like in Fig 6B, that the soma membrane current is the injected current minus current exiting the soma, so a lower *I*_m_ at *z* = 0 for *f* = 1000 Hz indicates more current exiting directly through the soma. **C:** Weighted average position of the amplitude of membrane currents (*λ*_AC_) above and below soma, and the *p*_*z*_ value, above and below soma at time *t*_max_.

The *p*_*z*_ amplitude from the pyramidal model decreased from 1.04 nAµm at 1 Hz to 0.046 nAµm at 1000 Hz. The *p*_*z*_ amplitude from the inhibitory L5 Martinotti cell instead decreased from 0.338 nAµm at 1 Hz to 0.0908 nAµm at 1000 Hz. The somatic membrane potential response of the L5 Martinotti cell would therefore only be a third of the L5 pyramidal cell response at 1 Hz, but twice as large at 1000 Hz.

The L4 bipolar cell has complete asymmetry around the soma (all dendrites are below the soma), and as such, we might expect it to exhibit the strongest tES response at all frequencies. At 1000 Hz, however, the L5 Martinotti cell has a somewhat higher amplitude (0.0815 versus 0.0908 nAµm) (Fig 1B). We found that this was caused by the quite different dendritic diameters of the cells: If both the L5 Martinotti cell and the L4 BP cell instead had the same uniform dendritic diameter (1 µm), the asymmetric L4 BP cell had a substantially larger *p*_*z*_ response at all frequencies (Sup Fig S4).

For the neuron with the smallest spatial extent (L5 NGC), the spatial distribution of membrane currents were very similar for 10 and 100 Hz (Fig 7B, bottom), resulting in very similar *p*_*z*_ amplitudes for the two frequencies.

To investigate the hypothesis further, we calculated *p*_*z*_ above and below soma for all neocortical neuron models (Fig 8). On average, the *p*_*z*_ asymmetry is substantially larger for pyramidal cells compared to inhibitory neurons at low frequencies. However, at high frequencies, much of the asymmetry is lost. In particular, *p*_*z*,above_ reduces strongly for pyramidal cells. Therefore, pyramidal cells and inhibitory neurons have a much more similar asymmetry in *p*_*z*_ at high frequencies.

**Fig 8.**
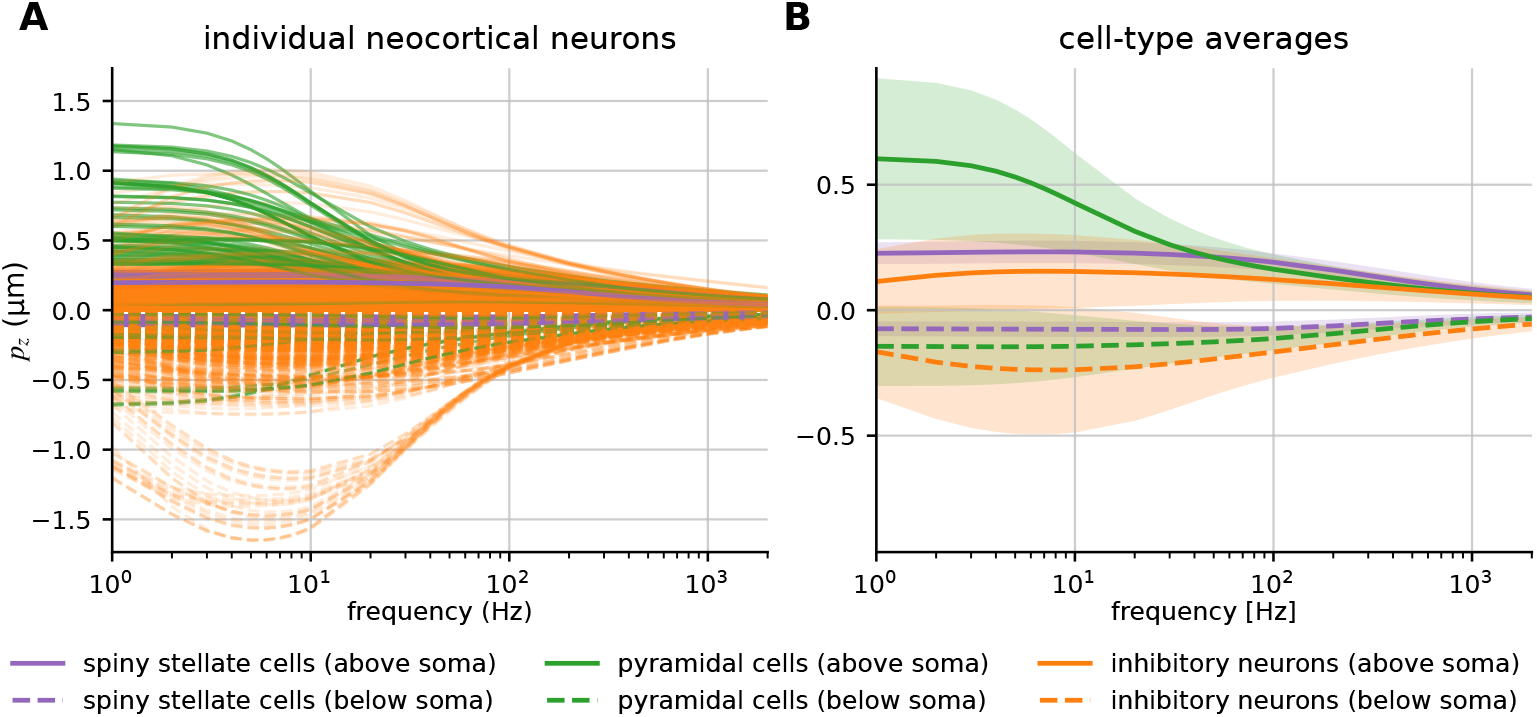
Pyramidal cells are, on average, effectively more asymmetric than interneurons at low frequencies but not high frequencies. **A:** *p*_*z*_ values above and below soma at the time *t*_max_ when the total *p*_*z*_ has its maximum value (see Methods). Most pyramidal cells have higher *p*_*z*_ above the soma compared to inhibitory neurons at low frequencies, while *p*_*z*_ below the soma is relatively small for most pyramidal cells, giving large asymmetry in membrane currents for pyramidal cells. *p*_*z*_ above the soma for pyramidal cells reduce markedly for increasing frequencies. **B:** The average *p*_*z*_ above and below soma for different neuron types. There is a marked reduction in average *p*_*z*_ above the soma of pyramidal cells with increasing frequencies.

### 2.4 Analytical formula for morphology-dependence of frequency response

The reciprocity theorem also enables us to find an analytical expression for a neuron’s response to transcranial electric stimulation (tES) for simplified neuron models. We first considered an idealized ball-and-two-sticks neuron consisting of a spherical soma with diameter *d*_s_, and two dendritic sticks with constant diameters *d*_d_ pointing in opposite directions along the *z*-axis. We derived a transfer function, **T**_*p*_, linking an arbitrary somatic input current **Î**_in_ to the resulting dipole moment in the *z*-direction 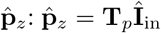. Here,

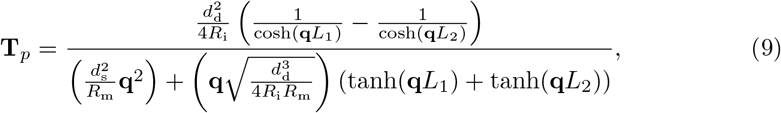

where 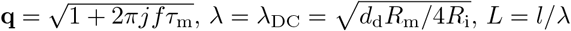, where *l* is the length of the dendritic stick, *d*_d_ is the diameter of the stick, and *d*_s_ is the soma diameter (see Sec S1 for derivation). This can be related to the amplitude of the membrane potential in the soma 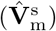 in response to tES via *k*,

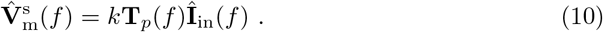

For the previously considered example location, an electric field of 1 mV/mm requires a tES current amplitude of 4.76 mA, which gives 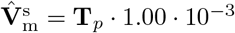 mV/µm.

The expression in the numerator of the transfer function in Eq (9) shows that a completely symmetrical neuron (*L*_1_ = *L*_2_) will generate no dipole moment from somatic current input, and therefore have no somatic membrane potential response to tES. More asymmetrical neurons, with a larger difference between *L*_1_ and *L*_2_ will, in general, have a higher response to tES. Further, Eq (9) shows that a larger dendritic diameter and a smaller somatic diameter give a larger soma *V*_m_ amplitude response to tES.

The effects of the total length (*L*_1_ + *L*_2_) and the frequency-dependency via **q** are not easily apparent from visual inspection of Eq (9). Therefore, we calculated 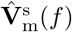 for a range of different ball-and-two-stick neurons across frequencies ranging from 1 to 2000 Hz (Fig 9). The longer neurons had higher 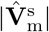 responses than shorter neurons at low frequencies (Fig 9A), but the response was more frequency-dependent, as also observed for the neocortical neurons (Fig 1). At 1000 Hz, the shorter neurons had higher 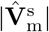 responses than the longer neurons. Furthermore, higher asymmetry, that is, higher relative difference between the lengths of the two sticks, gave clearly higher 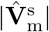 responses across all frequencies (Fig 9B). The soma and dendrite diameters (*d*_s_ and *d*_d_) had a smaller effect on the 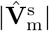 responses at low frequencies (Fig 9C and D), but the effect of *d*_d_ increased with frequency (Fig 9D).

**Fig 9.**
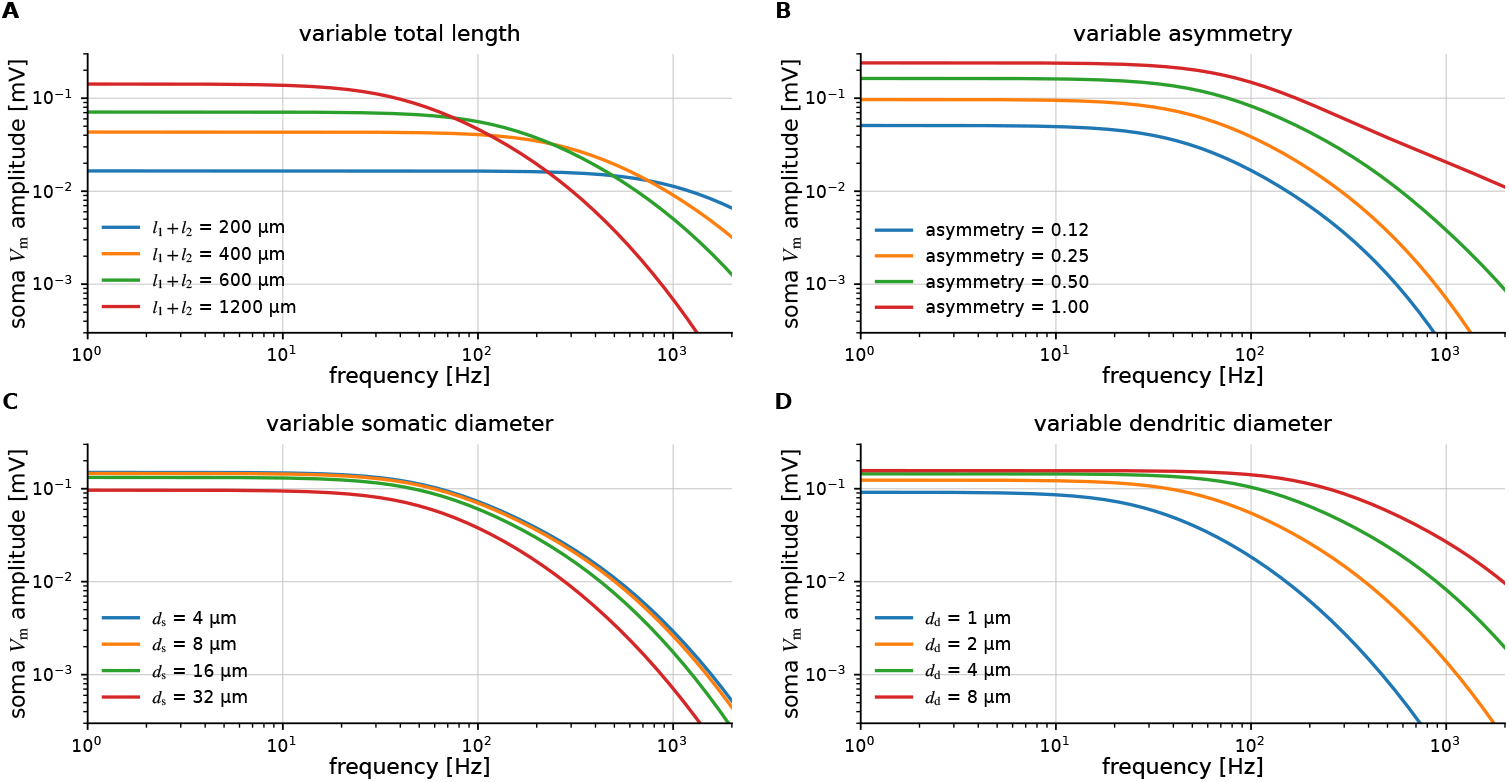
Analytic formula illustrates morphology dependence of tES response. Standard parameters: *l*_1_ = 670 µm, *l*_2_ = 335 µm, *d*_d_ = 2 µm, *d*_s_ = 20 µm and asymmetry-factor= 1*/*3 (*l*_2_ = *l*_1_*/*2) if the parameter is not varied. **A:** Varying the total length of the neuron, keeping asymmetry, soma diameter and dendrite diameter constant. **B:** Varying the asymmetry factor, defined as |*l*_1_ − *l*_2_ | */*(*l*_1_ + *l*_2_), keeping *l*_1_, dendrite diameter and soma diameter fixed. **C:** Varying the soma diameter, keeping lengths and dendrite diameter fixed. **D:** Varying the dendritic diameter, keeping lengths and somatic diameters fixed.

To further investigate the interplay of neuronal length and asymmetry, we calculated 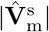 for different upper- and lower lengths (Fig 10) at *f* = 10 Hz and *f* = 1000 Hz. At *f* = 10 Hz, only very symmetrical neurons (analogous to L4 SS and L5 NGC) had low 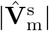 responses, and longer neurons had substantially larger responses (Fig 10A). The 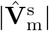 response increased quite uniformly with distance from the diagonal (which represents perfect symmetry, that is, no asymmetry). At *f* = 1000 Hz, however, only very asymmetrical neurons had large 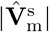 responses. The magnitude of 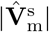 only increased with length up to about 250 µm (Fig 10B), after which the response decreased quite uniformly with distance from the edge (which represent perfect asymmetry). In other words, longer neurons need to have a much higher asymmetry in order to exhibit the same high 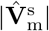 response at high frequencies.

**Fig 10.**
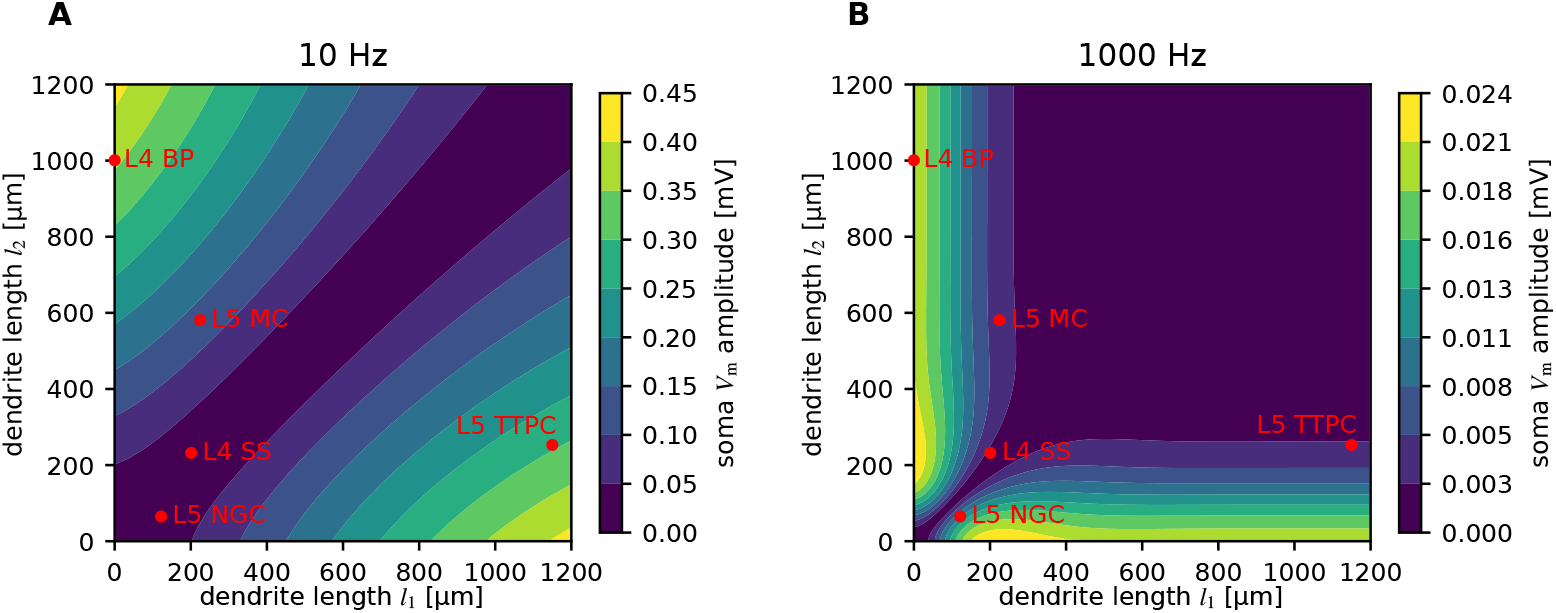
The effect of asymmetry at low and high frequencies. Contour plot of the 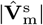 response for different dendritic lengths in the ball-and-two-sticks model at 10 Hz (**A**) and 1000 Hz (**B**). The soma diameter was *d*_s_=20 µm and dendrite diameters *d*_d_=2 µm. The lengths of the longest upwards-pointing (*l*_1_) and downwards-pointing (*l*_2_) dendrites are indicated with red dots for the five example cells (Fig 1A).

The ball-and-two-sticks model underlying the contour plot in Fig 10 assumes no branching and uniform passive parameters and dendritic diameters. The model is, therefore, too simplistic to quantitatively reproduce all the results from the biophysically detailed cell models. However, it can qualitatively explain many of the observed features of the cell-type specific frequency-dependent tES response from Fig 1B: (i) The L4 BP cell is both long and asymmetric, and is therefore strongly sensitive to tES for both low and high frequencies. (ii) The L5 TTPC cell is long and asymmetric enough to be sensitive to low-, but not high-frequency tES. (iii) The L5 NGC and L4 SS interneurons are both sufficiently symmetric to have weak responses at both frequencies. The L5 MC interneuron is of intermediate length and asymmetry, and the simple two-stick model would therefore predict an intermediate response as well. However, the two-stick model does not take into account the individual cell parameters, and it therefore is not able to predict the relatively strong high-frequency response of this cell model (Fig 1B, Fig S4).

The analytic expression for the ball-and-two-sticks in Eq (9) can be generalized to cell models with an arbitrary number of dendritic sticks protruding from the soma, with arbitrary diameters, passive parameters, and directions (see Methods, Eq 43). Such more complicated ball-and-sticks models can be used to mimic different morphological structures and cell types (Fig 11). From six different analytical example models, we again see that the completely asymmetric ball-and-stick models are most sensitive to tES across the frequency spectrum (black and gray).

**Fig 11.**
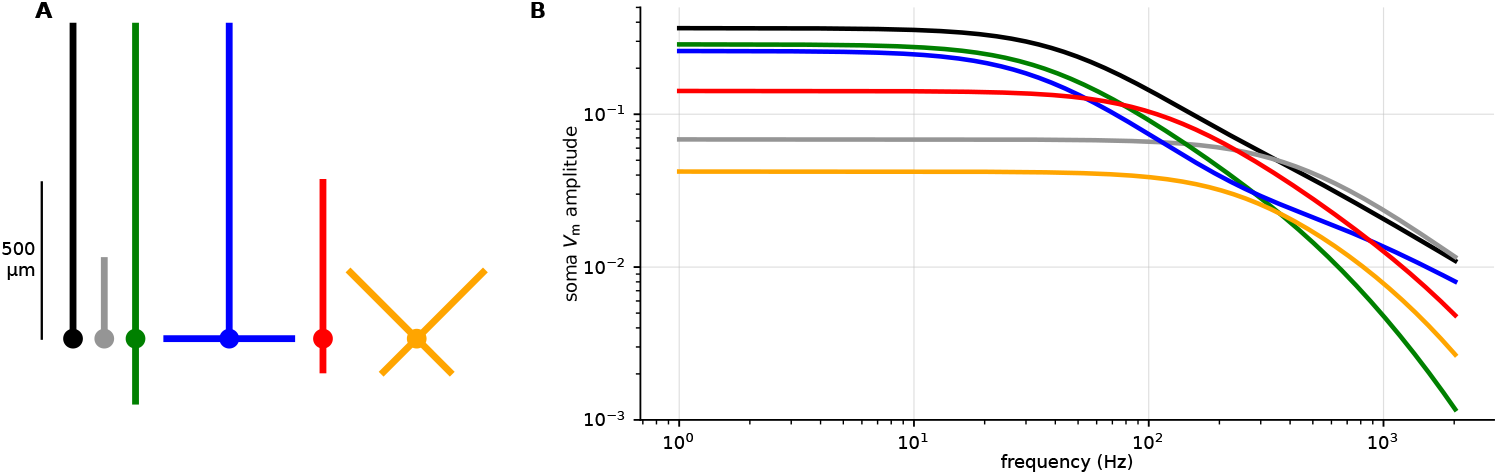
Analytic model of *V*_m_ responses for different types of morphologies. **A:** Six different simplifies morphologies including: (i) A classical ball-and-stick model with a single dendritic stick of length 1000 µm (black). (i) Same as in (i), but with a stick length of 250 µm (gray). (iii) A ball-and-two-sticks model where the sticks are oriented vertically but pointing in opposite direction (green). The upper stick length is 1000 µm while the lower stick length is 200 µm. (iv) Same as for (iii), except that there are two lower sticks that are oriented horizontally (blue). (v) A shorter version of the ball-and-two-sticks model from (iii) where the stick lengths are scaled down by a factor of two (red). (vi) A “classical” interneuron morphology with branches extending in different directions. To introduce some asymmetry, the upwards-pointing dendrites are 300 µm long, while the downwards-pointing dendrites are 150 µm long. For all models, the soma diameter was *d*_s_=20 µm and dendrite diameters *d*_d_=2 µm.

Interestingly, a short ball-and-stick model is actually slightly more sensitive to tES than the longer ball-and-stick model at frequencies above 300 Hz (Fig 11B, gray versus black). The reason is that, at high frequencies, there will be an increasing phase shift of the membrane currents along the stick, which cause cancellations, that is, membrane currents of opposite sign resulting in a smaller net current dipole moment. This effect can also be observed in Fig 10B.

A ball-and-two-sticks model where the sticks point in opposite directions had a similar low-frequency-response as the ball-and-stick model, but a substantially weaker high-frequency response (Fig 11, green), as also previously demonstrated (Fig 9). If the lower sticks instead pointed in the horizontal directions, the high-frequency response was more similar to the standard ball-and-stick model, and the cell model’s *V*_m_ response had a less steep decay with increasing frequency (Fig 11, blue).

Shorter ball-and-two-sticks models can have stronger high-frequency responses to tES than longer ball-and-two-sticks models (Fig 11B, red versus green), as also previously demonstrated (Figs 6 and 9).

Many types of interneurons have a high degree of symmetry around the soma, and as noted previously, perfectly symmetric cells will have negligible somatic tES responses. However, for small cells, even relatively modest asymmetries can result in higher tES sensitivity than for longer pyramidal-like cells (Fig 11B, orange versus green).

## 3 Discussion

In this study, we investigated cell-type specific responses to transcranial electric stimulation (tES) across a broad frequency range. In line with previous studies [20, 27–29], we demonstrated that pyramidal cells on average exhibited the strongest responses at low frequencies, whereas inhibitory neurons on average showed somewhat larger responses at higher frequencies (*f >* 60 Hz) (Fig 1). We showed that cell-type specific morphological properties can explain this (Figs 2, 3).

By leveraging the reciprocity theorem we demonstrated that the effect of tES on neurons is proportional to the same (frequency-dependent) current dipole moment that determines the contribution to the EEG signal (Fig 4), allowing us to investigate in detail how morphology affects neurons’ susceptibility to tES. In particular, we identified the key importance of dendritic asymmetry, that is, how spatially asymmetric dendrites are distributed around the soma along the axis of the imposed electric field.

We found that longer neurons are more sensitive to low-frequency tES—but also exhibit a more frequency-dependent tES response than shorter neurons—because of their longer frequency-dependent length constants (Fig 5). Most neurons have dendrites both above and below the soma, and in this case, longer neurons can effectively become more symmetric than shorter neurons at high frequencies (Figs 6, 7, 8). This helps explain why interneurons were more susceptible than pyramidal cells to high-frequency tES.

Finally, we derived an analytical formula for the somatic membrane potential response of idealized neuron models to an arbitrary tES. Using this formula, we extracted general insights into how morphological features affect how neurons respond to tES (Figs 9, 10, 11). This confirmed the importance of the dendritic asymmetry on the response to tES and demonstrated the counterintuitive insight that while total cell length increases tES sensitivity for low frequencies, it has the opposite effect at high frequencies (Figs 10, 11).

### 3.1 Implications of cell-type specific responses to electric Stimulation

These findings help us better understand the effect of tES at the fundamental level of single-cell membrane potential responses, and provide us with helpful rules-of-thumb. In particular, they demonstrate how principles governing neuronal responses to tES at low stimulation frequencies do not necessarily generalize to high-frequency regimes: While long neurons are more strongly affected by low-frequency stimulation, increased length can decrease sensitivity to tES at higher frequencies.

The reciprocity-based approach is excellently suited for implementing the effect of tES in point-neuron network models: It can take into account the full morphological complexity of a cell model and very accurately estimate the effect of the tES on the somatic membrane potential. It is then trivial to calculate what input current would be needed for the point neuron to experience this membrane potential on top of other ongoing activity. This approach can be expected to give more reliable results than network models of two-compartment cell models or ball-and-stick cell models, since it takes into account the effect of the entire detailed morphology. Note also that two-compartment models and ball-and-stick models are—in contrary to real pyramidal cells—perfectly asymmetric, which has a large effect on the expected tES sensitivity. At the same time, point neurons are more computationally efficient because of their simplicity.

These results suggest new opportunities for high-frequency tES by modulating inhibitory neuronal activity. Selective engagement of inhibitory activity may be advantageous for therapeutic applications targeting neurological disorders characterized by pathological hyperexcitability, including epileptic seizures [51] and Alzheimer’s disease [52]. Furthermore, reduced inhibition has been linked to depression [53, 54].

This study concerns single-cell effects of tES, but to understand tES at a functional level, it is necessary to study how tES affect networks. Simulations of tES at the network level typically uses a highly simplified model of the tES, like a fixed current input to point neurons or mean field models. This, however, does not take into account that the effect of tES depends on cell type and frequency. This study is therefore an important step towards a more accurate implementation of tES in network simulations.

### 3.2 Modelling assumptions

In our simulations, we applied the quasi-uniform approximation for the electric field, that is, stimulation of the neurons was performed with a spatially constant electric field across the size scale of individual neurons. In reality, the electric field caused by tES is not completely uniform, but the quasi-uniform approximation has been demonstrated to be quite accurate for tES [20, 38].

In the present study we have focused on the case where the applied electric field is oriented along the cortical depth direction (*z*-direction). The individual cell models in the database we used [36] were already aligned along the same axis, and no individual rotations were applied. Because of cortical folding, any given tES will, however, have different orientations at different cortical locations. For completeness, we therefore also simulated tES along the *x*- and *y*-axis on the neocortical neurons (Fig S7). On average, all different cell types showed higher soma *V*_m_ amplitudes responses to *z*-directed fields. Since the apical dendrites of pyramidal cells were oriented along the *z*-axis, these cells were not particularly strongly affected by the *x*- and *y*-directed fields at low frequencies, and along these axes, pyramidal cells and interneurons had relatively similar responses.

The cell models that we considered were based on experimental data [36]. We can therefore expect the morphological structures and diameters to be representative of real neurons, but we cannot exclude the possibility that passive parameters, like membrane resistances and axial resistances, included unintentional biases or errors. For example, many interneuron models showed a larger tES response at 10 Hz than at 1 Hz (Fig 1B). We found the reason for this to be that the somatic leak conductance was 10-fold larger than the dendritic leak conductance in these models (Fig S4). This 10-fold difference originates from the parameter optimization that was done in the construction of the models from patch-clamp recordings in [36]. It is, however, unclear how accurately these fitted values for the soma conductance reflect biology.

In this study, we only considered somatic membrane potential responses of neurons (sometimes referred to as the somatic doctrine [35]), the rationale being that the major effect from tES on neural activity comes from field-induced depolarization (or hyperpolarization) of the soma that affects the firing of action potentials. This might at first sight to be at odd with the expectation that the numerical change of membrane potential of axonal terminals from tES is larger than for somas [20, 35, 37, 55].

However, under *in vivo* conditions it can be expected that the tES induced action-potential firing relies on an effect called stochastic resonance [10] where the observed subthreshold fluctuations of the membrane potential in the somas set up by synaptic inputs are key. This synaptic noise may by itself bring the membrane potentials of some neurons to be so close to their firing threshold that a small sub-millivolt extra push from tES may modulate the firing.

Due to the lack of substantial synaptic input on the axons and axon terminals such a stochastic resonance effect is not expected here, arguing that it is the field-induced effect on the soma membrane potentials rather than the axonal membrane that drives the effects of tES in in vivo conditions, at least for the electric field magnitudes considered here. Note, however, that with axonal morphologies available, our modeling scheme (including the use of the reciprocity theorem), could if warranted equally well be used to calculate the tES responses in axons. It is, however, not clear that subthreshold polarizations of axonal terminals can be fully ignored, since such polarization has been reported to modulate neurotransmitter release [55].

Similarly, in this study, the effect of dendrites is only investigated in terms of how they affect the somatic membrane potential response. However, dendritic depolarization or hyperpolarization will also affect synaptic conductances by changing the driving force of synaptic inputs [16], and can affect the initiation of dendritic calcium spikes [56]. These questions warrants further attention.

### 3.3 Use of analytical formula

The analytical treatment offers a transparent way to examine how somatic membrane potential responses evolve with frequency, and how dendritic morphology above and below the soma shapes this dependence. Naturally, the simplified model cannot fully capture the structural complexity of morphologically realistic neurons, but since it allows arbitrary section-specific lengths, directions, diameters, and leak conductances, it is quite flexible. As an illustration of this, we adapted the parameters of the analytic formula to approximately mimic individual example cells (Fig S6).

The simplified model qualitatively reproduces cell-type specific frequency-dependent tES responses, helping explain why inhibitory neurons can become relatively more responsive than pyramidal cells at high frequencies (Fig 10). The formula is also computationally efficient, requires only a few lines of code, and relies on no special software libraries. It is, therefore, well suited for use together with network simulations of tES with point-neuron networks, or even mean-field models.

### 3.4 Application to networks

Conceptually the effect from electrical stimulation on the dynamics of a neural network can be thought of as a two-step process: First, each neuron is stimulated individually, then the effects from these individual stimulations affect the whole interconnected neural network. In this paper we have only modelled the first step, and in order to predict network effects these single-cell responses must incorporated in network simulations. Here we did not pursue this second step, but we believe the present work provides a key building block for pursuing such simulations.

To simulate such network effects, one must combine the present modelling of single-neuron stimulation with neural network models. Numerous such network models have been developed over the last decades and are ready to be used. For models based on biophysically detailed neuron models [57], results from using the present scheme for computing effects from transcranial electrical stimulation on single-neuron dynamics is directly applicable: (i) Compute the relevant component of the current dipole moment for each type of neuron in the network taking into account both the position and orientation relative to the direction to the stimulation electrode on the scalp. Here a detailed electric head model is required [47]. (ii) Use this computed current dipole moment to compute at each time step the addition to the soma membrane potential from the electrical stimulation for each neuron (see [37, their Fig 5]). Note that an analogous approach can be used to model network effects from intracranial electric stimulation, though here the current-dipole moment will in general not be sufficient to represent the effects from the different neuronal morphologies [37].

For networks based on point neurons such as integrate-and-fire neurons (e.g., [58–60], a hybrid approach, analogous to Hagen et al. (2016) [61], can be used where biophysically-detailed multicompartmental neurons first are used to calculate the effect of the electrical stimulation on the membrane potentials of each type of neuron. Next, such pre-computed membrane potentials can be added to the membrane potentials of the point neurons in the networks.

At present neural-field models based on population firing rates are the model type of choice for human whole-brain modeling using tools such as The Virtual Brain [62]. Kernel-based tools have been developed for biophysics-based prediction of electric brain signals generated by neuronal populations described by firing rates [63–65]. Similar approaches should be pursued to make kernel-based methods for predicting effects on population dynamics from electric stimulation.

The present study has highlighted the crucial dependence of neural responses to dendritic morphology, stimulation frequency and orientation of electric field relative to the neuron orientation. Further, given the mix of excitatory and inhibitory neurons in brain tissue, it is clear that it will be a challenge to make detailed prediction about the network-level effects pf various types of electric stimulation. However, given the present availability of (i) personalized detailed electric head model, (ii) techniques for computing single-cell responses to electric fields given neuronal morphologies as presented here, and (iii) the availability of biophysics-based network models, we have a good foundation for model-based exploration of effects of networks. Hopefully, this will contribute to make the clinical use of tES more accurate and reliable.

## 4 Methods

### 4.1 Simulation tools

All code was written in Python. The Python module LFPy 2.3 [66] was used for the simulations, running on NEURON 8.2.6 [67].

### 4.2 Neuron models

Neocortical neuron models were from the Blue Brain Project’s digital reconstruction of the neocortical microcircuit of juvenile rat [36].

When constructing the neocortical neurons with LFPy, the number of segments was increased to make sure that all segment lengths remained short enough at high frequencies. The neocortical neurons were made passive by removing all active conductances.

Idealized ball-and-sticks neurons were also constructed with LFPy. This idealized neuron model consist of a spherical soma with diameter *d*_s_, and two dendritic sticks with constant diameters *d*_d_ pointing in opposite directions along the *z*-axis, with identical neural parameters, i.e., same specific membrane resistance *R*_m_, specific membrane capacitance *C*_m_ and the inner resistivity *R*_i_.

### 4.3 Morphological properties

For the neocortical neuron models obtained from the Blue Brain Project, morphological data were extracted to investigate how neuronal morphology influences the response to electric stimulation. In particular, several length-related properties in the *z*-direction were stored: The distances from the soma to the upper and lower endpoints (respectively *l*_1_ and *l*_2_). We identify the endpoints by finding the compartment with the highest *z*-value, and the compartment with the lowest *z*-value. The total length in *z*-direction is then the sum of *l*_1_ and |*l*_2_|. An asymmetry-factor in the *z*-direction perpendicular to the cortical surface, was calculated as

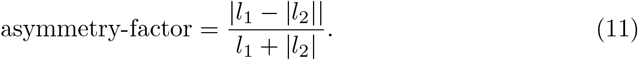

In addition, the soma diameter (*d*_s_) and a measure of the average dendrite diameter (*d*_d_) in the *z*-direction were found. The measure for average dendrite diameter was found by taking all compartments in the neuron primarily oriented in the *z*-direction (i.e. *dz > dx* and *dz > dy*) and finding the average diameter of these compartments.

### 4.4 Electric field stimulation

The neocortical neuron models from the Blue Brain Project’s digital reconstruction were stimulated with spatially constant extracellular electric fields with a strength of 1 mV/mm oriented in the *z*-direction along the cortical surface normal, mimicking transcranial electric stimulation (tES) with alternating current (tACS) with frequencies ranging from 1 to 2020 Hz.

The large and distant electrodes applying the current during transcranial electric stimulation (tES) creates an electric field in the brain with low spatial gradient on the size scale of individual neurons [20]. For this reason, we used the quasi-uniform assumption when modeling the effect of tES, meaning a negligible spatial gradient of the electric field on the local area of the neuron [38].

Extracellular stimulation was implemented using the `extracellular` mechanism provided by NEURON [68]. The extracellular potential was applied to the cell via the `insert_v_ext` cell class method from LFPy, which imposes the extracellular potential as a boundary condition [66].

To assess the neurons’ responsiveness to the stimulation the resulting membrane potential (*V*_m_) in the soma was recorded, and the amplitude of *V*_m_ was found via Fourier analysis (see Sec 4.6).

### 4.5 White noise current input

For the reciprocity-based analysis, the neuron models were stimulated with somatic white-noise current input (*I*_in_). The white-noise current was constructed as a sum of sinusoids, with frequencies ranging from 1 to 2020 Hz. Each frequency component had equal amplitude (5 pA unless otherwise specified) with a random phase [37, 48]. The current was applied as a point-process membrane current in NEURON [37].

From the white noise current stimulation, the resulting membrane currents (*I*_m_) across the neuron are recorded. From this, the *z*-component of the dipole moment (*p*_*z*_) was extracted by LFPy, and the amplitude of *p*_*z*_ across all frequency components were found through Fourier analysis (Sec 4.6).

The membrane currents used to calculate *λ*_AC_ are the amplitudes of the membrane currents in each compartment of the neurons, and also here the amplitudes are extracted via Fourier analysis (Sec 4.6).

### 4.6 Signal analysis

The Fourier analyses were performed using the *scipy.fftpack* Python package, structured with code from [69] to return frequency components and the corresponding amplitudes and phases of recorded signals during the simulations.

To relate each frequency component of the white-noise current stimulation to the resulting membrane currents and dipole moments, we used the phases obtained from the Fourier analysis to evaluate these quantities at specific times. Because the input current, membrane currents, and dipole moments can each have different phases—and because the dipole moments above and below the soma may also differ from the total dipole moment—we reconstructed a sinusoid for each frequency component using its amplitude and phase.

### 4.7 Feature importance

Linear regression analysis was performed using scikit-learn’s `LinearRegression` function, which fits the model by minimizing the sum of squared errors. Model performance was evaluated using scikit-learn’s `r2_score` function to compute the *R*^2^ value. Prior to regression, predictor variables were standardized using scikit-learn’s `StandardScaler` . Feature importance values were defined as the resulting regression coefficients.

### 4.8 Frequency-dependent length constant

The frequency-dependent length constant *λ*_AC_ used, is derived in Pettersen and Einevoll (2008) [50] as

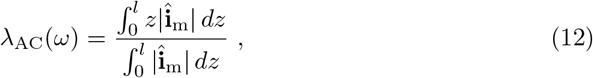

for a sinusoidally varying membrane potential 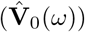, imposed as a boundary condition at the soma end (*z* = 0) of a stick with a sealed and at *z* = *l*. With complex notation (bold), the membrane current density for a single frequency component *f* is given by 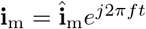. The transmembrane currents can be written as

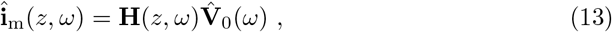

where

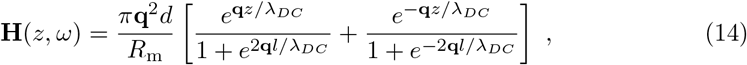

and 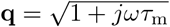 introduces the complex number, where *τ*_m_ = *R*_m_*C*_m_ is the membrane time constant, from the specific membrane resistance (*R*_m_) and the specific membrane capacitance (*C*_m_). We simplified *λ*_AC_ to

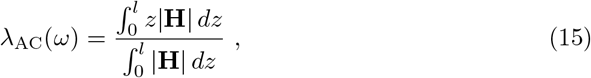

that was evaluated numerically in Python using 1000 spatial sample points along the stick and the trapezoidal integration rule.

### 4.9 Analytical formula

The transfer function, **T**_*p*_ relating a somatic input current **Î**_in_ and the resulting current dipole moment in the *z*-direction 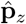 is given by Pettersen et al. [70] for a ball-and-stick neuron. Based on this, **T**_*p*_ for the ball-and-two-sticks neuron model (see Sec 4.2) is given by (see Sec S1)

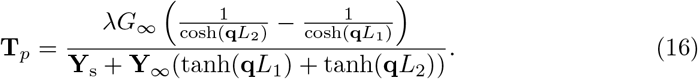

where *L* = *l/λ* and *l* is the length of the dendritic stick, 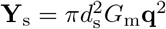 is the soma admittance and **Y**_∞_ = **q***G*_∞_ is the admittance of an infinite dendritic stick, where 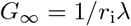 is the infinite-stick conductance. Here, the dependency of frequency is not introduced via *λ*_AC_. Instead, we use the typical definition 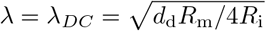, and the frequency-dependence is introduced via 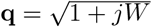, where *W* = *ωτ*_m_ is the dimensionless frequency.

With these parameters written out, Eq 16 can be written,

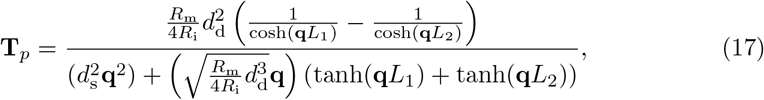

where *d*_s_ and *d*_d_ are the soma and dendrite diameters, respectively. For more details, see Sec S1.

The transfer function was related to the amplitude of the membrane potential in the soma 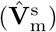 via *k*,

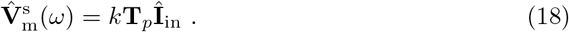

The transfer function and the theoretical 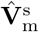 were implemented in Python, and the absolute value was calculated.

To confirm the theoretical calculation of 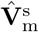, it was compared with numerical simulations of ball-and-sticks neurons, recording the amplitudes of the membrane potential *V*_m_ in the soma resulting from electric field stimulation (Fig S5).

## 4.10 Code Availability

The data used in this work can be reproduced by the simulation code, available at https://github.com/SusanneDahle/CellTypeDependenceElStim.git.

## A Supporting information

### S1 Derivation of the transfer function for ball-and-sticks models

This derivation is based on the work in Pettersen et al. (2014) [70].

#### Complex cable equation

We start with the cable equation for a cylinder with a constant diameter *d*_d_, which is given by

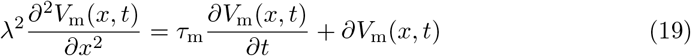

where we have the neuron length constant 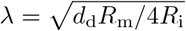 and the membrane time constant *τ*_m_ = *R*_m_*C*_m_. These constant indicate the temporal and spatial duration of an electric signal in the cable. The membrane potential *V*_m_ falls with 1*/e* after time *τ*_m_ or after the length *λ* [21, p.58], depending on how much current that exits the cell through the membrane. This is decided by the electrical properties of the neuron: The specific membrane resistance (*R*_m_), specific membrane capacitance (*C*_m_) and inner resistivity (*R*_i_), the specific membrane conductance, *G*_m_ = 1*/R*_m_ will also be used. With dimensionless variables the cable equation can be written as

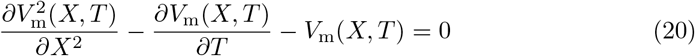

where *X* = *x/λ* is the dimensionless length and *T* = *t/τ*_m_ is the dimensionless time.

Our goal is to understand a neuron’s response to different frequencies of an input current. Because the cable equation is linear in the subthreshold domain, each frequency-component of the input signal can be can be treated individually via Fourier. Therefore it is useful to use complex notation (bold text) where *V*_m_ is expressed in the frequency domain as

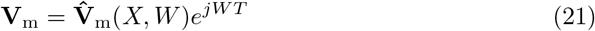

where *W* = *ωτ*_m_ is the dimensionless frequency (*ω* = 2*πf*). Now 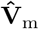 contains both the amplitude and phase of the membrane potential, and is related to *V*_m_(*X, T* ) via the Fourier components of the potential, given by

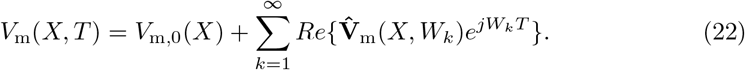

If we perform the Fourier transform, using 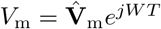 and considering a single frequency component *W*_*k*_, this can be substituted in to Eq. 20, and we get the cable equation in complex notation, simplified to

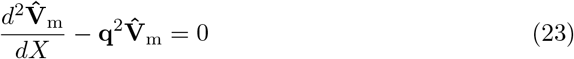

where **q**^2^ ≡ 1 + *jW* . For a finite stick with the electrotonic length *L* = *l/λ*, where *l* is the stick length, the general solution Eq. 23 can be written as

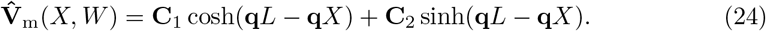

#### Specific Solution for a dendritic stick

In order to find the specific solution to the cable equation for a dendritic stick (cylinder) attached to a soma receiving an input current, we will use the boundaries with input at *X* = 0 and sealed end at *X* = *L*. When there is a current input in the soma, the input current creates an initial membrane potential 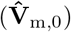 in the soma which acts as the driving force for the current flow though the dendritic stick. At *X* = 0, where the dendrite is attached to the soma, the membrane potential is continuous, which gives the first boundary condition 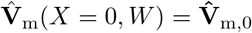. The sealed end boundary condition means no axial current though the end of the stick. The axial current in the dendritic stick is given by

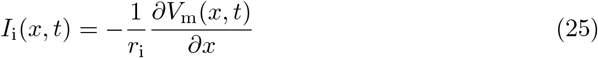

where 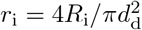 is the inner resistance per unit length of cable. With complex notation and dimensionless variables this is written as

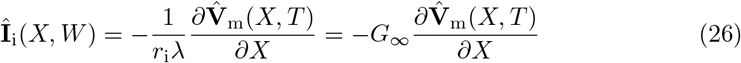

where *G*_∞_ = 1*/r*_i_*λ* is the infinite-stick conductance. The second boundary condition is then 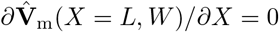. These two boundary conditions gives the specific solution to the cable equation for the dendritic stick as

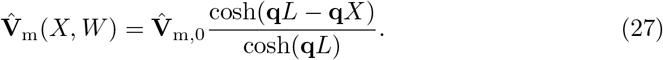

#### Input admittance

The initial membrane potential 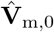 can be related to the input current **Î**_in_ via the input admittance 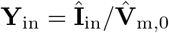. If we first consider a ball-and-stick neuron receiving an input current **Î**_in_ in the soma, due to Kirchhoff’s current law, **Î**_in_ has to be the equal to the sum of current leaving the soma through the membrane, i.e., the somatic membrane current **Î**_m,s_, and the current leaving the soma into the dendritic stick, i.e.,the axial current in the dendrite **Î**_i_(*X, W* ) at *X* = 0. The soma can be represented as an RC-circuit, which with complex notation gives the somatic membrane current density 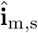 as

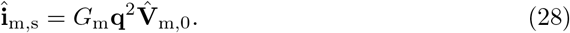

A spherical soma with diameter *d*_s_ gives the area of the soma 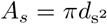, and the input admittance 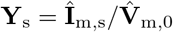 will therefore be given as

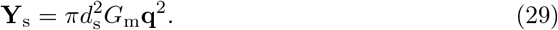

The current leaving the soma going into the dendrite is given by axial current **Î**_i_(*X, W* ) at *X* = 0. By inserting the specific solution to 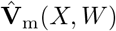 from Eq. 27 into the equation for the axial current in Eq. 26 we get

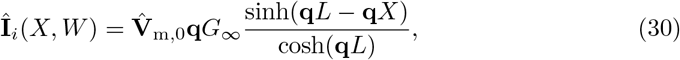

which gives the input admittance of the dendritic stick 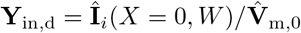 as

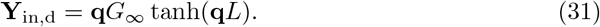

For an infinite stick lim_*L*→∞_ tanh(**q***L*) → 1, the infinite-stick admittance **Y**_∞_(*W* ) = **q***G*_∞_.

By Kirchoff’s current law, the total input admittance for the ball-and-stick is given by **Y**_in_ = **Y**_s_ + **Y**_d_. If the neuron instead consist of *N* number of dendritic sticks, the input current will spread to all *N* sticks, and the input admittance is therefore now affected by the sum of all individual dendritic stick admittances. This gives

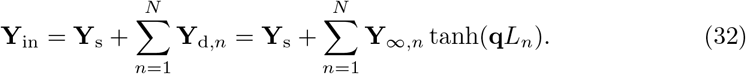

#### Dipole moment from a dendritic stick

In order to find an expression for the dipole moment from somatic current input we must look at the membrane currents through the dendritic stick. This is given as

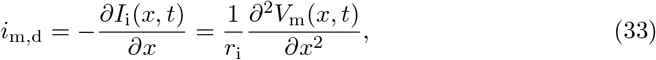

which with complex notation is expressed as

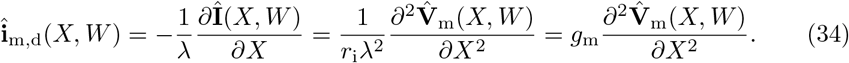

With the specific solution for 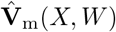 in Eq. 27, we get the membrane current in the dendritic stick as

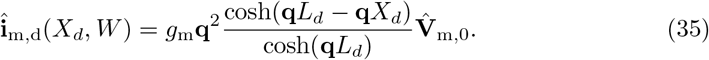

The dipole moment is the sum of all membrane currents multiplied with their position (Eq. 4). With a continuous cable, the complex dipole moment from the dendritic stick with a sealed end is given by the integral

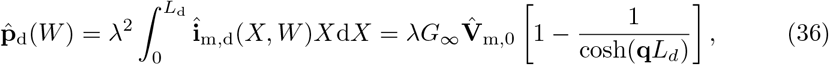

and have the direction of the dendritic stick.

#### Transfer function

Our goal is to find a transfer function relating the dipole moment in the *z*-direction 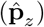 and the input current **Î**_in_ in the soma as

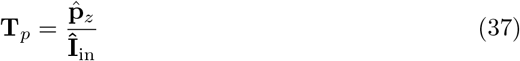

The dipole moment from a dendritic stick 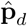, given in Eq. 36 has the direction of the stick. This dipole moment projection onto the *z*-axis is further given by 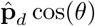 where *θ* is the angle between the direction of the stick and the *z*-direction. For *N* number of dendritic stick, the total dipole moment in the *z*-direction 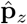 is then

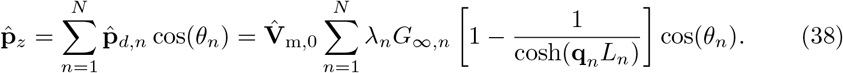

This gives the transfer function

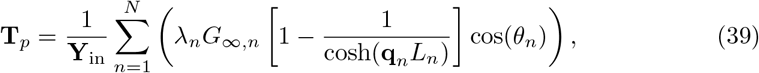

since 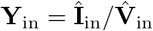.

#### Transfer function for Ball-and-Two-Sticks

We consider the case where the neuron has two dendritic sticks point in opposite directions along the *z*-axis and have identical neural parameters and identical dendrite diameter *d*_d_, we then have

- **Directions:** *θ*_1_ = 0^°^ =⇒ cos *θ*_1_ = 1, and *θ*_2_ = 180^°^ =⇒ cos *θ*_2_ = −1.
- **Parameters:** *λ*_1_ = *λ*_2_, *G*_∞,1_ = *G*_∞,2_, *Y*_∞,1_ = *Y*_∞,2_, **q**_1_ = **q**_2_.

The transfer function can then be simplified to

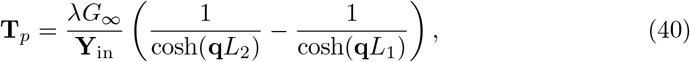

where the total input admittance is:

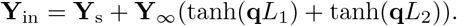

This full expression for the transfer function Ball-and-Two-Sticks neuron is then

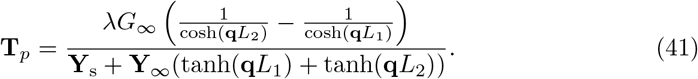

To gain insight into how different neural geometries affect the transfer function, 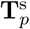 can be written out as

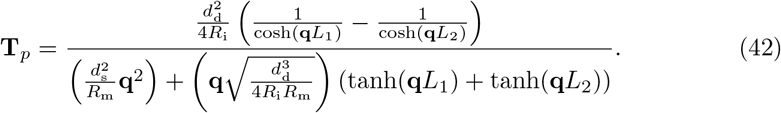

Following the same approach, this can be generalized to an arbitrary number of dendrites with individual lengths, diameters, and angles *θ*_*i*_. This gives,

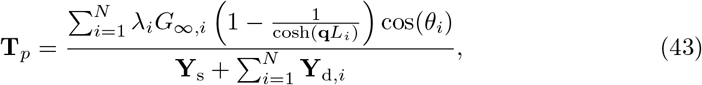

where the different components are listed in Tab S1.

**Table S1.** List of symbols and parameters in alphabetical order.

| Symbol | Default value (Unit) | Description |
| --- | --- | --- |
| $C_m$ | 0.01 pF/$\mu\text{m}^2$ | specific membrane capacitance |
| $d_d$ | 2 $\mu\text{m}$ | dendrite diameter |
| $d_s$ | 20 $\mu\text{m}$ | soma diameter |
| $f$ | 1 – 2000 Hz | frequency |
| $G_m = 1/R_m$ | 0.333 pS/$\mu\text{m}^2$ | specific membrane conductance |
| $G_\infty = \pi d_d^2/4R_i\lambda$ | 2.09 nS | infinite-stick conductance |
| $L = l/\lambda$ | (1) | electrotonic length |
| $l$ | ($\mu\text{m}$) | stick length |
| $\mathbf{q} = \sqrt{1 + jW}$ | (1) | frequency dependence |
| $R_i$ | 1.5 M$\Omega\mu\text{m}$ | inner resistivity |
| $R_m$ | 3 M$\Omega\mu\text{m}^2$ | specific membrane resistance |
| $T = t/\tau_m$ | (1) | dimensionless time |
| $W = \omega\tau_m$ | (1) | dimensionless frequency |
| $X = x/\lambda$ | (1) | dimensionless position |
| $\mathbf{Y}_{\text{in}}$ | (S) | input admittance |
| $\mathbf{Y}_s = \pi d_s^2 G_m \mathbf{q}^2$ | (S) | soma admittance |
| $\mathbf{Y}_d = \mathbf{q} G_\infty \tanh(\mathbf{q}L)$ | (S) | dendrite admittance |
| $\mathbf{Y}_\infty = \mathbf{q} G_\infty$ | (S) | infinite-stick admittance |
| $\lambda = \sqrt{d_d R_m/4R_i}$ | 1 mm | neuron length constant (DC) |
| $\tau_m = R_m C_m$ | 30 ms | membrane time constant |
| $\omega = 2\pi f$ | (rad/s) | angular frequency |

Selection of relevant symbols and parameters for the transfer function explaining dipole moment per input current. Table adapted from [70].

### S2 Supplementary figures and tables

**Fig S1.**
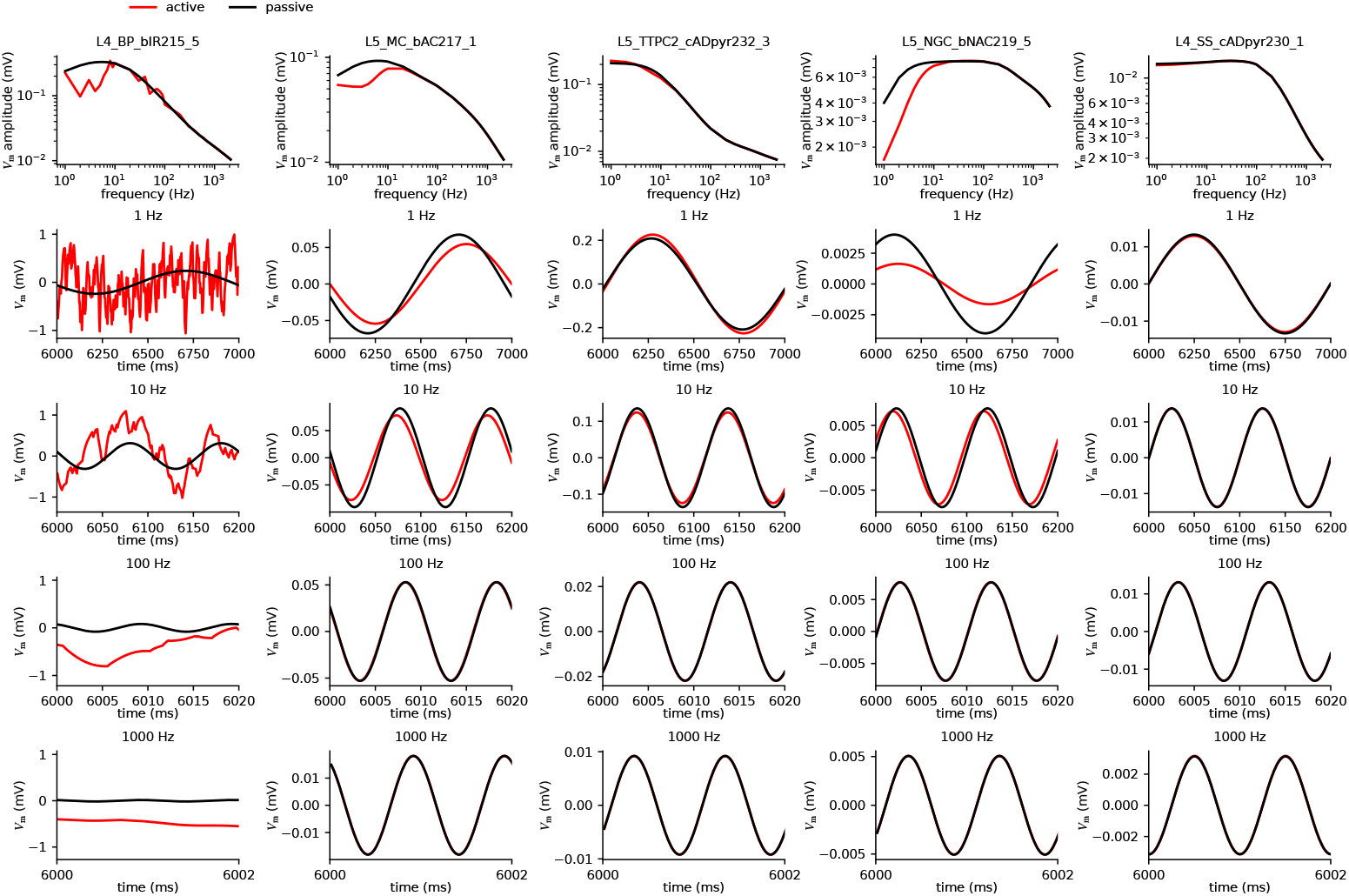
Comparison of active and passive cell models. Column correspond to different cell models. The first row is the somatic *V*_m_ response as a function of frequency, and the rows below are examples in the time-domain for different chosen frequencies (1 Hz, 10 Hz, 100 Hz, 1000 Hz). The cell model named “L4 BP bIR215 5” (leftmost column) has a stochastic potassium channel (“StochKv.mod”) which explains the “noise” in the active model.

**Fig S2.**
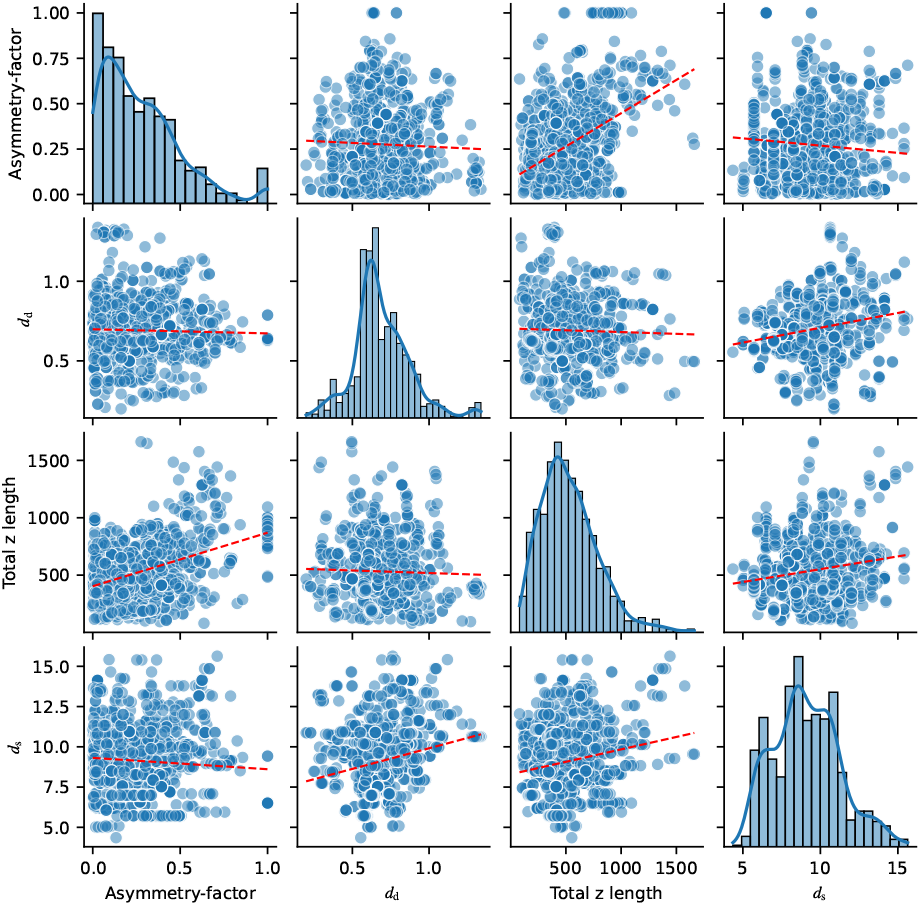
Relationship between selected properties for feature importance analysis. The selected properties are total *z* length, symmetry factor, average compartment diameter, and somatic diameter. Most features are largely independent, but the symmetry factor and total length have some correlation. Dendritic diameter and asymmetry factor calculations are explained in Sec 4.3.

**Fig S3.**
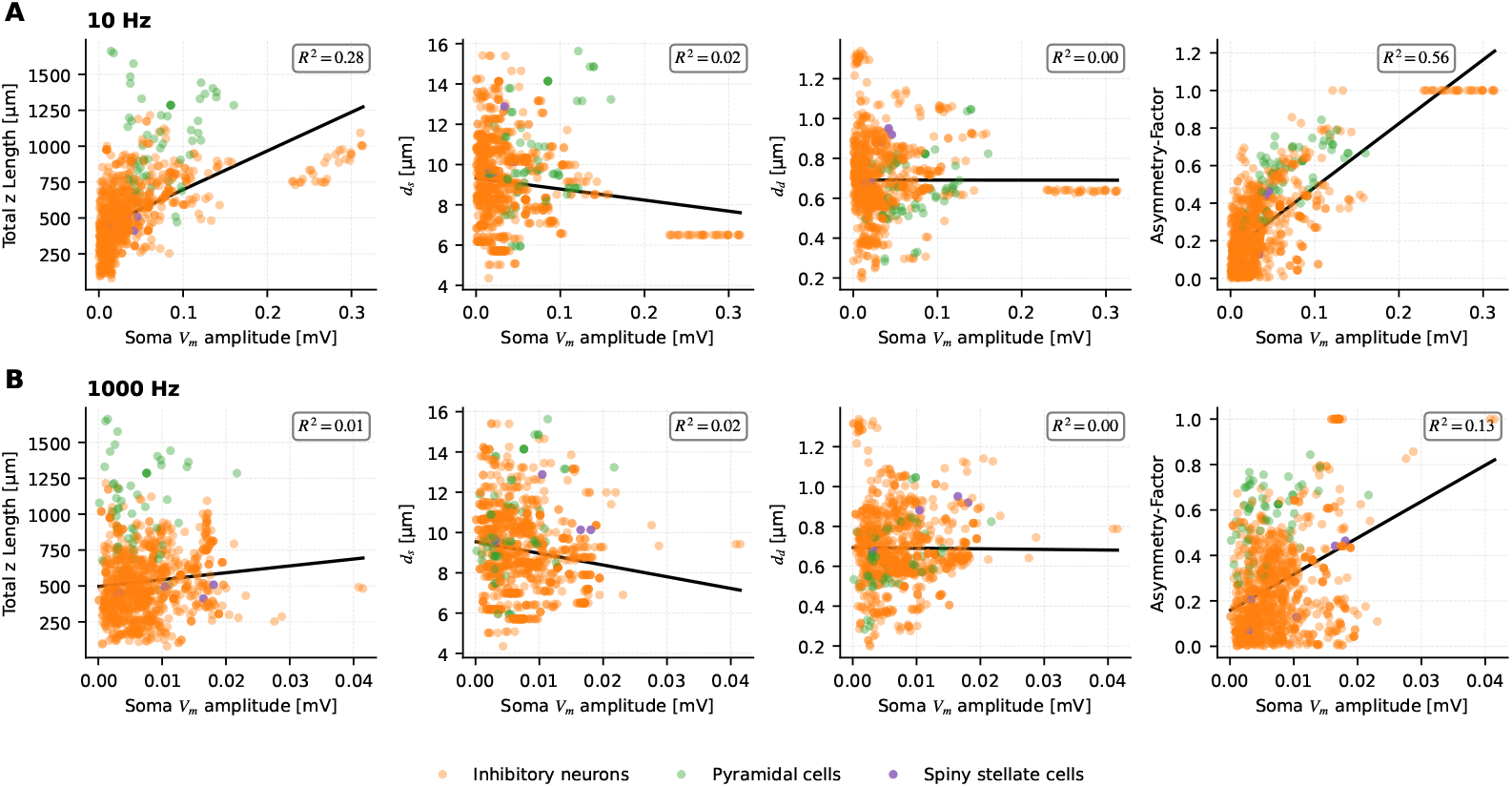
Relation between morphological properties in neocortical neurons and soma *V*_m_ amplitude in response to external electric field stimulation with A: *f* = 10 and B: 1000 Hz. *R*^2^-scores from single-feature regression models using the morphological property to predict soma *V*_m_ amplitude response. The asymmetry factor have the strongest correlation to soma *V*_m_ amplitude at both frequencies with *R*^2^ = 0.56 10 Hz at and *R*^2^ = 0.13 at 1000 Hz. All properties have lower (or the same) *R*^2^ scores at *f* = 1000 Hz, especially total *z* length. Asymmetry factor and *d*_d_ calculations are explained in Sec 4.3).

**Fig S4.**
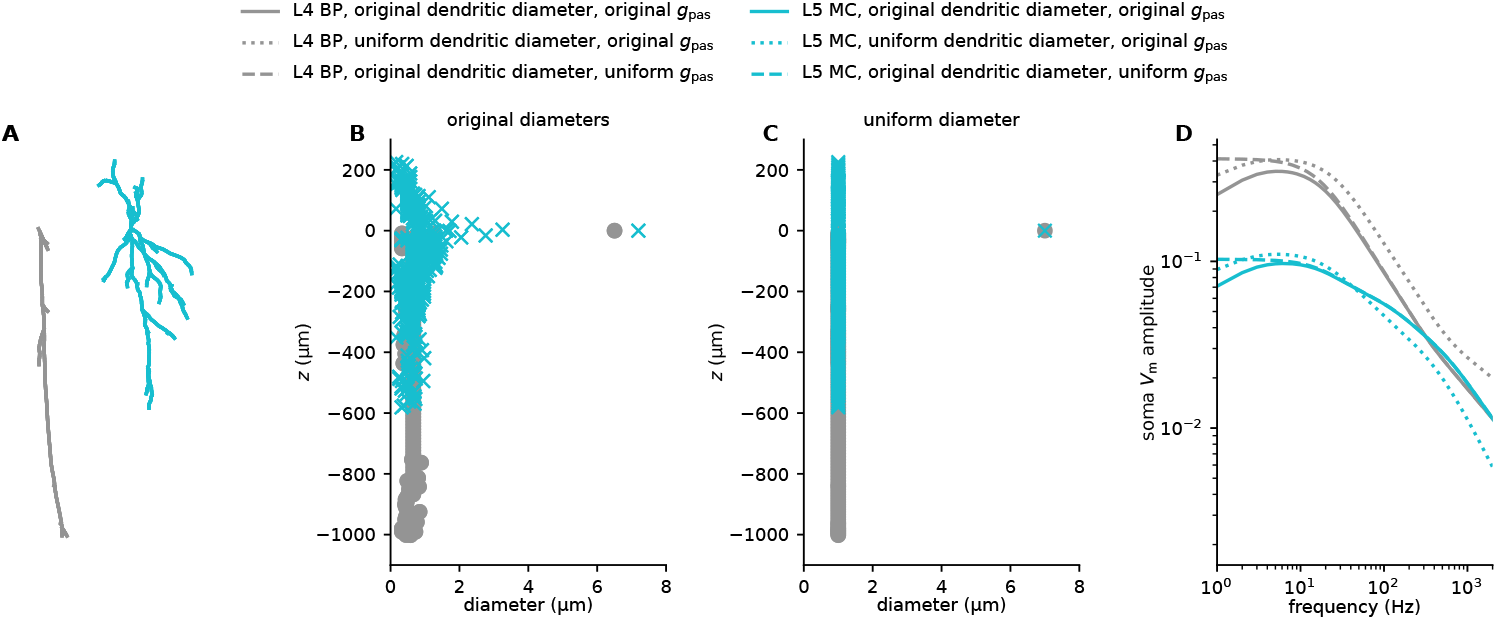
The effect of spatially variable parameters on soma *V*_m_ response. For two example interneuron models (L4 BP and L5 MC), we demonstrate the effect of spatially heterogeneous properties. The L4 BP cell is perfectly asymmetric (all dendrites are on the same side of the soma), and we might, therefore, naively expect it to be more sensitive to tES across the frequency spectrum than the less asymmetric L5 MC cell. However, at 1000 Hz the L5 MC cell is slightly more sensitive (see main text regarding Fig 10B). This is because of the effect of the individual diameters, and if both cell models had the same uniform dendritic diameters, the L4 BP cell would indeed be more sensitive to tES across the frequency spectrum (panel D, gray versus cyan dotted lines). A second example of the effect of spatially heterogeneous properties is that many interneurons models were more sensitive to tES at 10 Hz than at 1 Hz (see Discussion, Sec 3.2), and we found this to be caused by the fact that many interneurons had a 10-fold higher passive leak conductance in the soma than in the dendrites. For a uniform passive leak conductance, this resonance-like peak in the frequency spectrum disappeared (panel D, gray versus cyan dashed lines).

**Fig S5.**
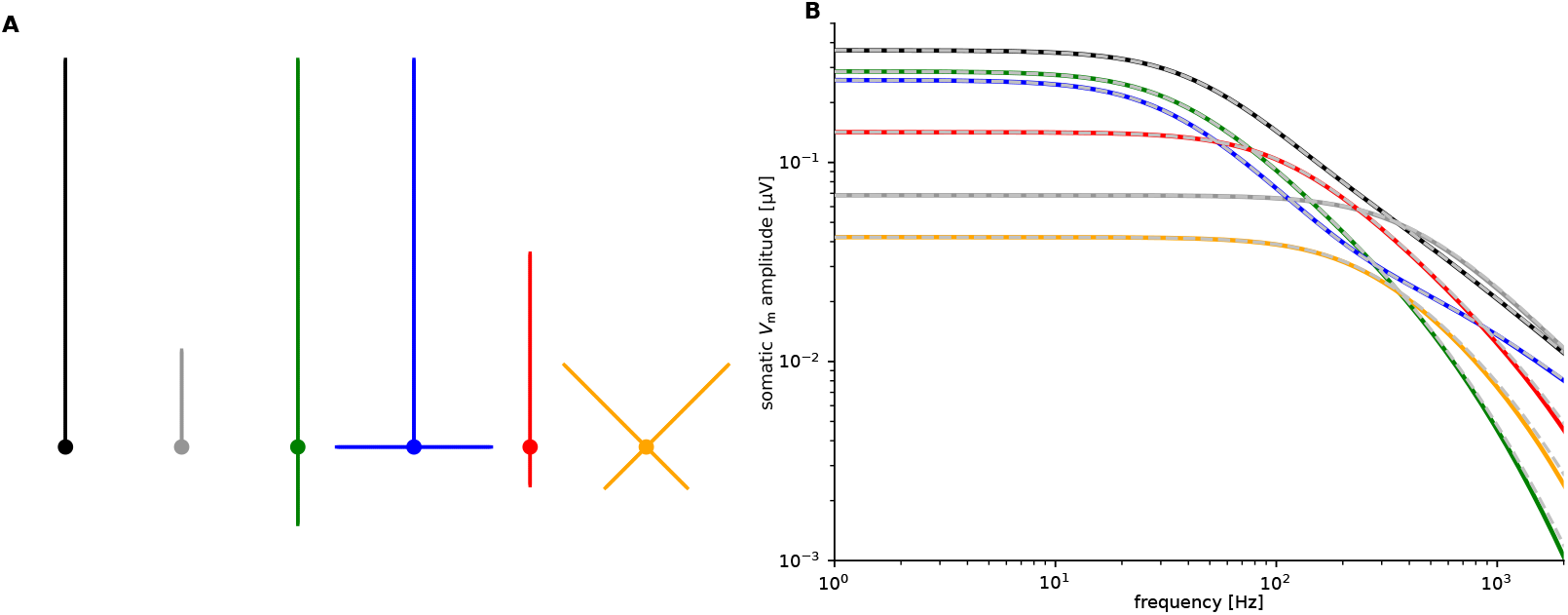
Validation of analytical formula. Calculated 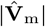 for different ball-and-sticks neuron models (different colors in panel A). Each model was simulated numerically in LFPy under the influence of an external electric field, and their resulting soma *V*_m_ amplitudes were recorded (full colored lines in panel B). These numerical results are compared to predictions from the analytic model (gray dotted lines in panel B), and the analytical calculations are able to predict the soma *V*_m_ amplitudes of the ball-and-sticks models stimulated with an external electric field.

**Fig S6.**
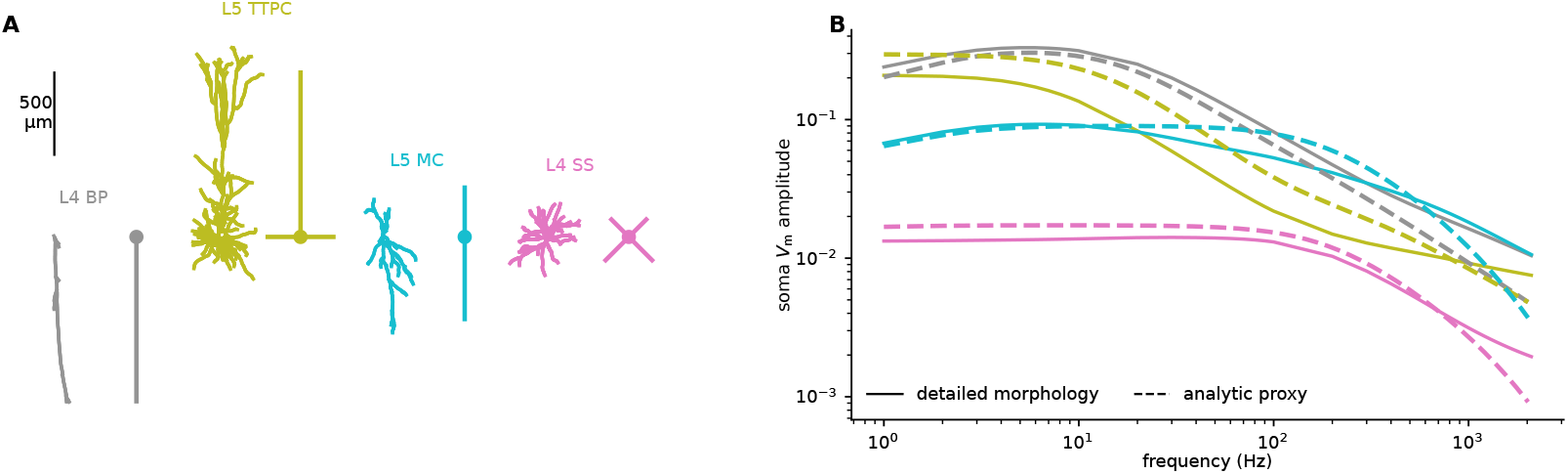
Comparison between example morphologically detailed cells and similar analytical models. **A:** Examples of morphologically detailed cells and simple analytic approximations (different colors). **B:** Somatic membrane potential responses of both the detailed cell models (full lines with colors corresponding to cells in panel A), and the analytic approximations (dashed lines with colors corresponding to cells in panel A). Note that only a course hand tuning was used on the analytic models. Applying numerical optimization techniques would certainly lead to a better fit, but the objective here is just to illustrate that the diverse effect of morphology on tES sensitivity can be approximately represented by the analytic formula.

**Fig S7.**
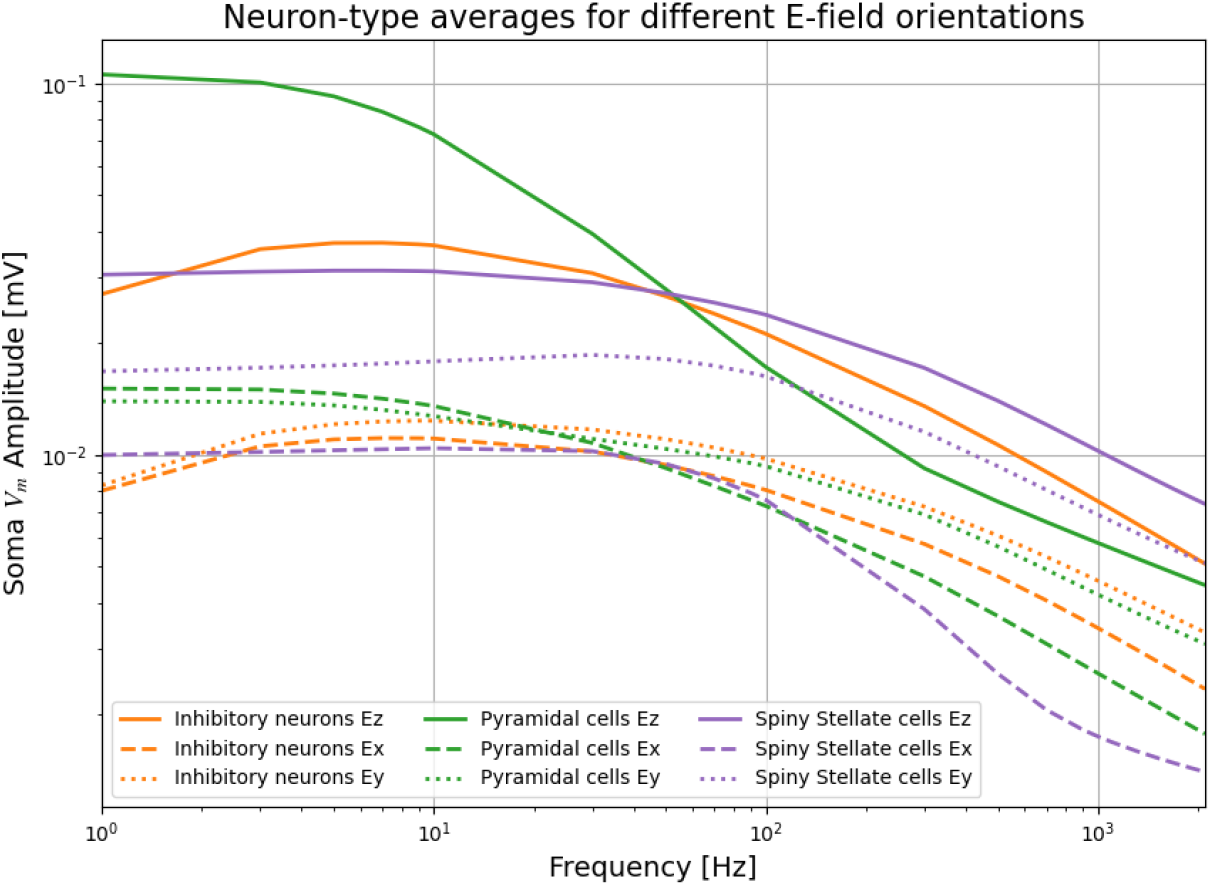
Average soma *V*_m_ amplitude response of neocortical cell types from external electric fields stimulation with different orientations. Cell-type averages from *z*-oriented electric field are the same as in Fig. 1D. The *z*-oriented electric field stimulation gives highest average soma *V*_m_ amplitude for all cell types in the neocortex. For all orientations, pyramidal cells have higher average soma *V*_m_ amplitude than inhibitory neurons at low stimulation frequencies, while inhibitory neurons have a higher average soma *V*_m_ amplitude than pyramidal cells at high stimulation frequencies (*f >* 60 Hz). The most pronounced difference in response from different orientations of the electric field is observed for pyramidal cells at low frequencies, while responses are more comparable at high frequencies. A one sided Mann-Whitney U test comparing soma *V*_m_ amplitudes between inhibitory neurons and pyramidal cells at *f* = 1000 Hz gave *p* = 0.00267 for *x*-directed fields, and *p* = 0.173 in *y*-directed fields.

**Fig S8.**
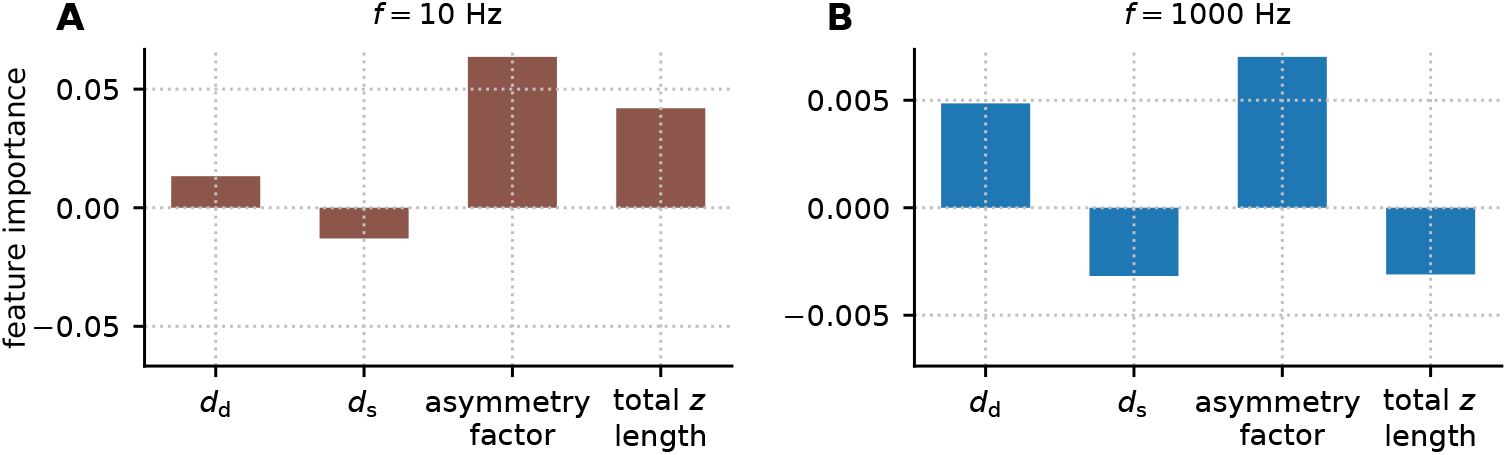
Feature importance values from linear regression on idealized ball-and-two-sticks models. Linear regression model included four selected properties to predict soma *V*_m_ amplitudes of ball-and-two-sticks models, resulting from applied electric field oriented in the *z*-direction perpendicular to the cortical surface. **A:** At *f* = 10 Hz, the regression model achieved *R*^2^=0.88. The asymmetry factor and total length had the highest absolute feature importance values. **B:** At *f* = 1000 Hz, the regression model achieved *R*^2^=0.77. The asymmetry factor still had the highest absolute feature importance value, while total length had the opposite effect as for 10 Hz. The morphological parameters were independently and randomly chosen with uniform probability density within certain ranges. The total length was in the range 200-1000 µm, the asymmetry factor was between 0 and 1, the somatic diameter was in the range 5-20 µm, and the dendritic diameter was in the range 0.5-2 µm. In total, 1000 cell models were included.

